# Two independent approaches converge to the cloning of a new *Leptosphaeria maculans* avirulence effector gene, *AvrLmS-Lep2*

**DOI:** 10.1101/2020.10.02.322479

**Authors:** Ting Xiang Neik, Kaveh Ghanbarnia, Bénédicte Ollivier, Armin Scheben, Anita Severn-Ellis, Nicholas J. Larkan, Parham Haddadi, W.G. Dilantha Fernando, Thierry Rouxel, Jacqueline Batley, Hossein M. Borhan, Marie-Hélène Balesdent

## Abstract

*Leptosphaeria maculans*, the causal agent of blackleg disease, interacts with *Brassica napus* (oilseed rape, canola) in a gene-for-gene manner. The avirulence genes *AvrLmS and AvrLep2* were described to be perceived by the resistance genes *RlmS* and *LepR2*, respectively, present in the cultivar Surpass 400. Here we report cloning of *AvrLmS* and *AvrLep2* using two independent methods. *AvrLmS* was cloned using combined *in vitro* crossing between avirulent and virulent isolates with sequencing of DNA bulks from avirulent or virulent progeny (Bulked-Segregant-Sequencing) to rapidly identify one candidate avirulence gene present in the effector repertoire of *L. maculans. AvrLep2* was cloned using a bi-parental cross of avirulent and virulent *L. maculans* isolates and a classical map-based cloning approach. Taking these two approaches independently, we found that *AvrLmS* and *AvrLep2* are the same gene. Complementation of virulent isolates with this gene confirmed its role in inducing resistance on Surpass 400 and Topas-*LepR2*. The gene renamed *AvrLmS-Lep2* encodes for a small cysteine-rich protein of unknown function with an N-terminal secretory signal peptide, which are common features of the majority of effectors from extracellular fungal plant pathogens. The *AvrLmS-Lep2* / *LepR2* interaction phenotype was found to vary from a typical hypersensitive response to intermediate resistance sometimes at the edge of, or evolving toward, susceptibility depending on the inoculation conditions. *AvrLmS-Lep2* was nevertheless sufficient to significantly reduce the stem lesion size on plant genotypes with *LepR2*, indicating the potential efficiency of this resistance to control the disease in the field.

## INTRODUCTION

Stem canker, or blackleg disease, caused by the ascomycete *Leptosphaeria maculans*, is a major disease of oilseed rape (canola, *Brassica napus*) (Fitt *et al*., 2006). Gene-for-gene interactions between *L. maculans* and *B. napus* have been described. At least 18 resistance (*R*) genes have been reported for the *L. maculans-Brassica* interaction, with only two cloned to date; *LepR3* and *Rlm2* (Delourme *et al*., 2004; Yu *et al*., 2005; Delourme *et al*., 2006; Rimmer, 2006; Yu *et al*., 2008; Van de Wouw *et al*., 2009; Long *et al*., 2011; Larkan *et al*., 2013, 2015). In contrast, eight avirulence (*Avr*) genes have already been cloned from *L. maculans*; *AvrLm1, AvrLm2, AvrLm3, AvrLm4-7, AvrLm5-9, AvrLm6, AvrLm10* and *AvrLm11* (Gout *et al*., 2006; Fudal *et al*., 2007; Parlange *et al*., 2009; Balesdent *et al*., 2013; Van de Wouw *et al*., 2014; Ghanbarnia *et al*., 2015; Plissonneau *et al*., 2016, Ghanbarnia *et al*., 2018, Petit-Houdenot *et al*., 2019). Map-based cloning of the first *AvrLm* genes (*AvrLm1, AvrLm6* and *AvrLm4-7;* Gout *et al*., 2006; Fudal *et al*., 2007; Parlange *et al*., 2009) took many years due to the lack of a *L. maculans* genome sequence and their location in the repeat-rich, gene-poor regions of the genome (Rouxel *et al*., 2011). More recently, cloning of *AvrLm11, AvrLm2* and *AvrLm5-9* showed that the availability of a reference genome and a repertoire of effector genes facilitated the identification of candidate avirulence genes (Balesdent *et al*., 2013; Ghanbarnia *et al*., 2015; 2018).

The use of race-specific resistance genes in Brassica species (*Rlm* and *LepR*) is an efficient way to control stem canker. However, such resistances can be rapidly overcome following emergence and selection of virulent isolates in fungal populations (i.e., Sprague *et al*., 2006). Four *R* genes (*LepR1, LepR2, LepR3* and *LepR4*) were genetically characterised in *Brassica rapa* subsp. *sylvestris* (Yu *et al*., 2005, 2008, 2013). *B. rapa* subsp. *sylvestris* was used as a source of resistance to *L. maculans* in the 1990s (Crouch *et al*., 1994) and introduced into *B. napus* cv. as “*sylvestris-*derived resistance”. From 2000 to 2003, the *B. napus* variety Surpass 400 and derivative cultivars containing *sylvestris-*derived resistance were grown on large acreages across Australia before the resistance was overcome in the Eyre Peninsula region of South Australia (Li *et al*., 2004; Sprague *et al*., 2006). Based on the genetic analysis of fungal isolates that overcame the Surpass 400 resistance, Van de Wouw *et al*. (2009) reported that at least two avirulence genes, *AvrLm1* and *AvrLmS*, conveyed avirulence towards Surpass 400, supporting the idea of an *RlmS-AvrLmS* interaction in the host plant. Genetic analyses of the Surpass 400 resistance described either one or two resistance loci (Li & Cowling, 2003; Yu *et al*., 2008; Long *et al*., 2011), termed *LepR3* (Yu *et al*., 2008), BRLMR1 and BRLMR2 (Long *et al*., 2011), or *LepR3* and *RlmS* (Larkan *et al*., 2013). Larkan *et al*. (2013) mapped and cloned the resistance gene *LepR3* from Surpass 400 and showed that *LepR3* recognizes the AvrLm1 protein. *LepR3* can therefore be considered a functional homologue of *Rlm1*, though *LepR3* and *Rlm1* reside on different chromosomes (A10 and A07, respectively). It was also demonstrated that a second resistance gene was present in Surpass 400, likely corresponding to the *AvrLmS* avirulence gene, i.e. *RlmS* (Larkan *et al*., 2013). However, uncertainties on the *RlmS* resistance remain since Yu *et al*. (2008) noted the possible presence of *LepR2* or a similar gene in Surpass 400 along with *LepR3*.

Dissection of *R* gene content in a variety and conclusions on genetic determination of the resistance only rely on phenotypic evaluation of the interaction based on inoculation tests with *L. maculans* field isolates, whose avirulence (AVR) gene content may differ from one study to the other. Due to epistatic effects between avirulence genes, a single gene control of a resistant phenotype toward a given isolate may hide a more complex *R* gene determination due to the lack of adequate differential isolates. Cloning of the matching AVR genes can thus help us understand the genetic basis and potential equivalence, or at least functional redundancy, between *R* genes with different names. In the current study, we report on the cloning of the avirulence gene corresponding to *RlmS* and *LepR2* using two independent approaches. On the one hand, the AVR gene recognized by *LepR2, AvrLep2*, was cloned following a standard map-based cloning approach, whilst, the gene interacting with *RlmS, AvrLmS*, was cloned using a bulked-segregant sequencing (BSS) strategy. The two approaches identified the same avirulence gene, which shares all characteristics of *L. maculans* avirulence effector genes. Noticeably, the two strategies were efficient although the resistant interaction phenotype appeared to be variable depending on environmental conditions. This work also suggests that *LepR2* and *RlmS* may be the same Brassica *R* gene. Until that is determined we will refer to the common effector as *AvrLmS-Lep2*.

## RESULTS

### Approach 1: Bulked segregant sequencing (BSS)

#### Phenotypic characterization of X82 progeny for bulked segregant sequencing

A progeny was created for BSS, following a cross (#82) between isolates WT50 and INV13.269, segregating for avirulence on *B. napus* cv. Surpass 400. Parental and progeny isolates phenotyping on an extended *B. nap*us differential set revealed that *AvrLm6, AvrLm4-7* and *AvrLmS* segregated in cross #82 (Table 1). Their segregation fitted the expected 50:50 ratio (Table 1). In addition, eight phenotypic classes were recovered in the progeny, with ratios fitting the hypothesis of independence between *AvrLm6, AvrLm4-7* and *AvrLmS* (Table 1, *X*^*2*^=8.25, *P*=0.689) as previously established (Van de Wouw *et al*., 2009). Following this phenotyping step, 47 isolates were selected for the preparation of six bulks of DNA. Bulk 1 contained DNA from 25 progeny isolates with an avirulent phenotype on Surpass 400 but either virulent or avirulent on *Rlm6* or *Rlm7;* Bulk 2 comprised DNA from 22 progeny isolates virulent on Surpass 400; Bulk 3 contained DNA from 24 avirulent isolates on *Rlm7*, but either virulent or avirulent on *Rlm6* or *RlmS* and Bulk 4 contained DNA from 23 isolates virulent on *Rlm7*; Bulks 5 and 6 contained isolates being avirulent on *RlmS* and *Rlm7*, or virulent on both genes, respectively (Table 2).

**Table 1.**
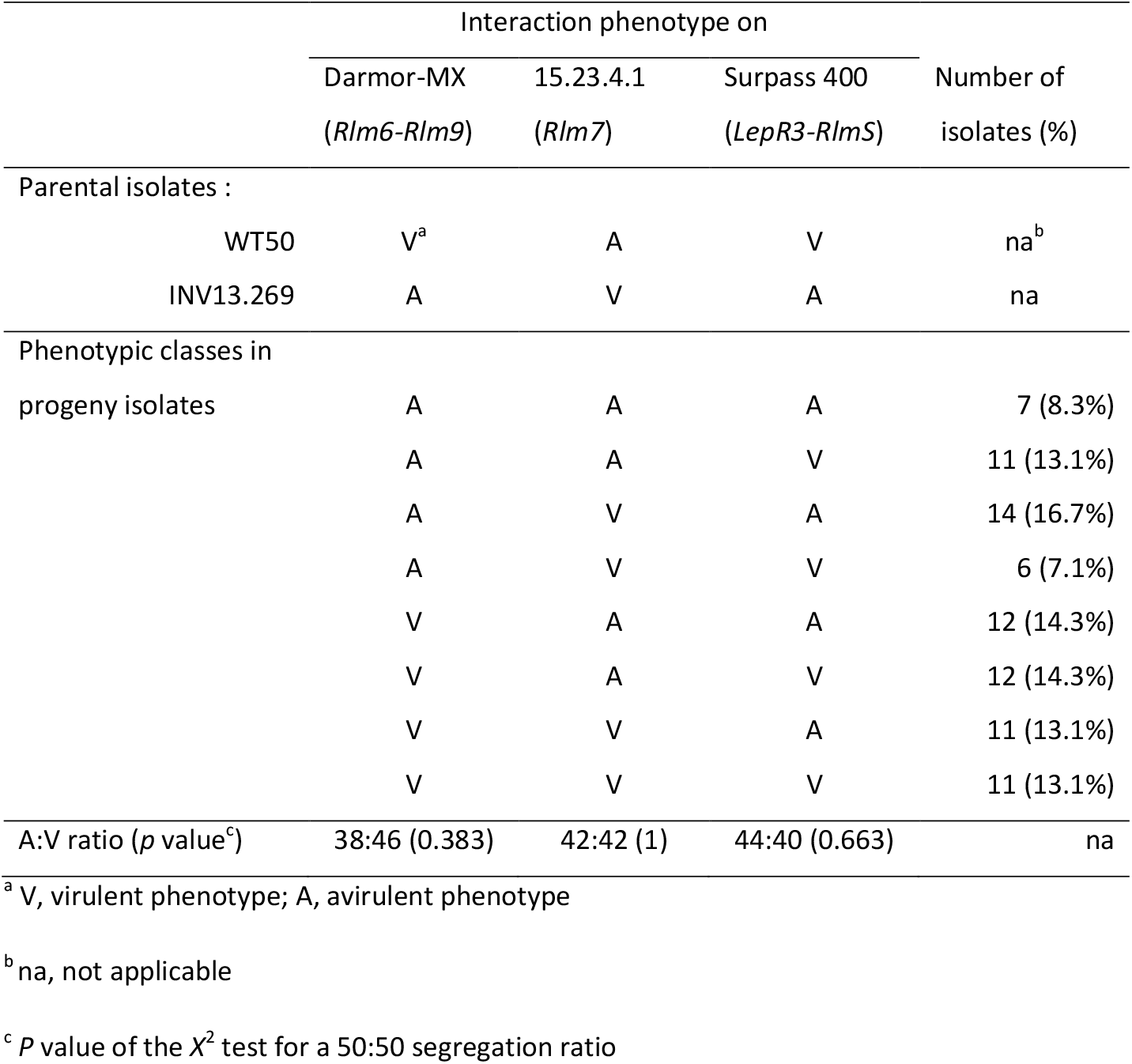
Segregation for virulence on *Rlm6, Rlm7* and *RlmS* (Surpass 400) in the *Leptosphaeria maculans* cross X82 (WT50 x INV13.269)

**Table 2.** Characteristics of DNA bulks and sequence mapping results.

| Bulk or sample No. | Bulk name (abbreviated name) | Type of isolate | No. of contributing isolates in the bulk | Nb of reads mapped to Lmb_jn3_08343 | Fraction of Lmb_jn3_08343 bases covered by at least one read |
| --- | --- | --- | --- | --- | --- |
| 1 | *AvrLmS* (AS) | Progeny, avirulent on Surpass 400 | 25 | 626 | 1.00 |
| 2 | *avrLmS* (aS) | Progeny, virulent on Surpass 400 | 22 | 2 | 0.37 |
| 3 | *AvrLm7* (A7) | Progeny, avirulent on *Rlm7* | 24 | nd | nd |
| 4 | *avrLm7* (a7) | Progeny, virulent on *Rlm7* | 23 | nd | nd |
| 5 | *AvrLmS* + *AvrLm7* (AS7) | Progeny, avirulent on Surpass 400 and *Rlm7* | 11 | 809 | 1.00 |
| 6 | *avrLmS* + *avrLm7* (aS7) | Progeny, virulent on Surpass 400 and *Rlm7* | 9 | 0 | 0.00 |
| 7 | INV13.269 (a7AS) | Parental isolate | 1 | 1056 | 1.00 |
| 8 | WT50 (A7aS) | Parental isolate | 1 | 1 | 0.03 |

#### BSS statistics and validation of the BSS strategy

The Illumina whole-genome sequencing generated reads (2 x 150 bp) from the six bulks and the two parental isolates. After quality trimming, the average number of reads generated was in the range of 47 – 75 million and the average genome coverage depth ranged between 164 X and 272 X (Supplementary Table 1). In total, 65,727 SNPs were identified between the parental isolates, excluding 19,650 SNPs in repetitive regions. The average number of SNPs found in the bulked progeny was 64,473 (Supplementary Table 2). After quality filtering for QTL mapping in the bulked pairs, the total number of SNPs retained was 27,532 (Bulks 3/4), 27,128 (Bulks 1/2) and 26,010 (Bulks 5/6).

To validate the BSS strategy for AVR gene cloning, a QTL-Seq analysis was carried out using Bulk 3 (A7) vs. Bulk 4 (a7), differing for the gene *AvrLm4-7*. A QTL was found on scaffold JN3_SC03 (genome version of Dutreux *et al*., 2018) at position 62,390 to 345,050 (Supplementary Figure 1, Supplementary Tables 3-4). This region contains 50 predicted genes (Lmb_jn3_03239 to Lmb_jn3_03288) including *AvrLm4-7* (GenBank nucleotide sequence AM998638.1, Lmb_jn3_03262). This result validates the BSS strategy for identification of a genomic region containing a gene of interest and suggests that the size of the bulks (23 and 24 isolates) is adequate for this purpose.

**Table 3.** Pathogenicity test for *L. maculans* isolates (wild-type and transformants) on *B. napus* lines carrying diverse blackleg *R* genes.

| Isolates/Transformants<sup>a</sup> | *B. napus* lines/cultivars and R gene content |  |  |  |  |  |  |  |  |  |  |
| --- | --- | --- | --- | --- | --- | --- | --- | --- | --- | --- | --- |
|  | Topas (T) | T-*Rlm1* | T-*Rlm2* | T-*Rlm3* | T-*Rlm4* | Vulcan | Roxet | Goéland | T-*LepR1* | T-*LepR2* | T-*LepR3* |
|  | *Control* | *Rlm1* | *Rlm2* | *Rlm3* | *Rlm4* | *Rlm6* | *Rlm7* | *Rlm9* | *LepR1* | *LepR2* | *LepR3* |
| v23.1.3 | V<sup>b</sup> | A | V | V | A | A | A | V | A | V | A |
| 00-100 | V | V | A | A | V | A | V | A | A | A | V |
| v23.1.3: *AvrLep2* (AW1) | V | A | V | V | A | A | A | V | A | A | A |
| v23.1.3: *AvrLep2* (AW2) | V | A | V | V | A | A | A | V | A | A | A |

<sup>a</sup>v23.1.3: *AvrLep2* (AW1) is a transformant with pNL11-*AvrLep2* construct (*AvrLep2* avirulent allele was amplified from isolate 00-100 with native promotor);
v23.1.3: *AvrLep2* (AW2) is a transformant with pLM4-*AvrLep2* construct (*AvrLep2* avirulent allele coding region was amplified from isolate 00-100, driven by
*AvrLm1* promoter.
<sup>b</sup> Interactions classified as either virulent (V) or avirulent (A).

**FIGURE 1.**
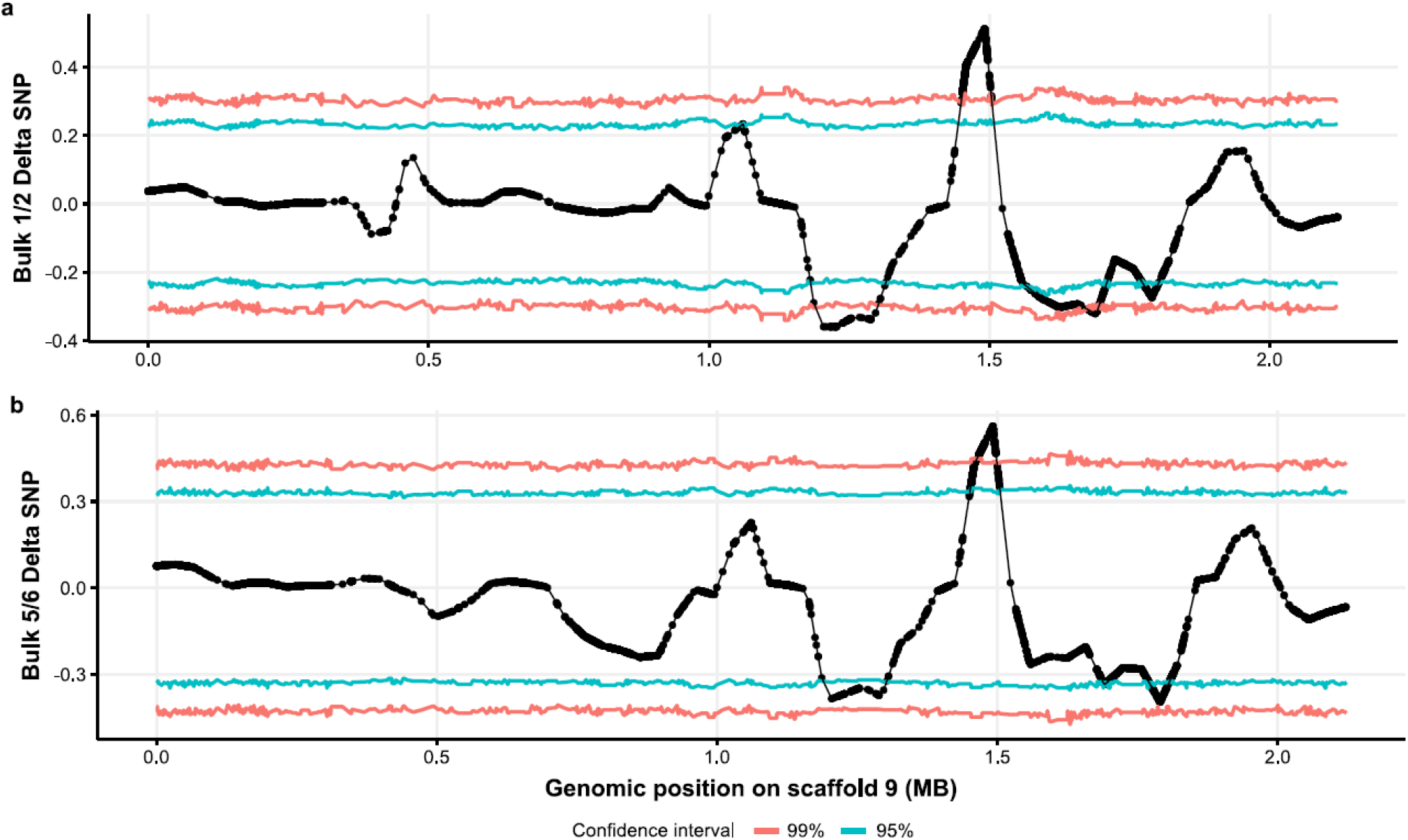
Plot of Δ(SNP-index) between Bulks differing for *AvrLmS* across scaffold JN3_SC09. (a) Bulks 1/2 (*AvrLmS*) and (b) Bulks 5/6 (*AvrLmS, AvrLm7*). Confidence intervals of 95% (red) and 99% (blue) are shown. QTL coordinates are provided in Supplementary Tables 5 and 6.

#### Identification of a candidate gene for *AvrLmS* using BSS

QTL-Seq results for Bulk 1 (AS) vs. Bulk 2 (aS) and Bulk 5 (AS7) vs. Bulk 6 (aS7) were compared to identify the candidate region for *AvrLmS*. The analysis revealed a QTL for *AvrLmS* within a 335 kb (Bulks 5/6) or 816 kb (Bulks 1/2) region on JN3_SC09 (Figure 1, Figure 2, Supplementary Tables 4-6). The QTL for both bulked pairs overlapped, with the major QTL supported by the most SNPs and the highest ΔSNP value showing peaks at position 1,481,733 in Bulks 1/2 and Bulks 5/6. Candidate SNP analysis for *AvrLmS* using Bulk 1/2, Bulk 5/6 and both parents (Samples 7/8) confirmed the QTL-Seq results, identifying a total of 437 genome-wide SNPs that segregate with the avirulence trait. Of these, 410 were found on JN3_SC09, with all SNPs found in a 398 kb region overlapping with the QTL region (position: 1,477,092-1,874,868 bp). This region contains 28 genes (Lmb_jn3_08331 to Lmb_jn3_08358). The candidate region also contains a 285 kb AT-rich region (JN3_SC09:1,533,065-1,818,564), enriched in repeats (Supplementary Figures 2-3, Supplementary Table 7), typical for genomic regions encompassing AVR genes in *L. maculans* (Rouxel *et al*., 2011). Only one gene, Lmb_jn3_08343, was located in this AT-rich region. The number of reads mapped to Lmb_jn3_08343 was over 600 for each of the *AvrLmS* bulks and the *AvrLmS* parent INV13.269, with every single base of the coding sequence covered by reads (Table 2). Comparatively, only three reads mapped to Lmb_jn3_08343 in the *avrLmS* bulk sample and the virulent parent WT50 (Table 2). Read coverage analysis revealed a ∼3 kb region (1,611,953-1,614,969 bp), with zero or close to zero base coverage in all the *avrLmS* bulk samples (Bulk 2, Bulk 6 and Parent WT50) whereas all the *AvrLmS* bulk samples (Bulk 1, Bulk 5 and Parent INV13.269) had per base coverage between 100 to 400 within this region. The ∼3 kb putative deletion contains the candidate gene Lmb_jn3_08343 (Figure 2, Supplementary Figure 3). This gene fulfils all criteria for a *L. maculans* avirulence gene candidate: (i) sequence variation, either in terms of SNPs or presence/absence variation (ii) genomic location in a gene-poor, AT-rich region and (iii) lack of sequence homology with other AVR genes or any protein in the database, except a weak homology with another candidate effector gene in *L. maculans* (Lmb_jn3_03815; 39.50% identity, e-value=1e-20).

**FIGURE 2.**
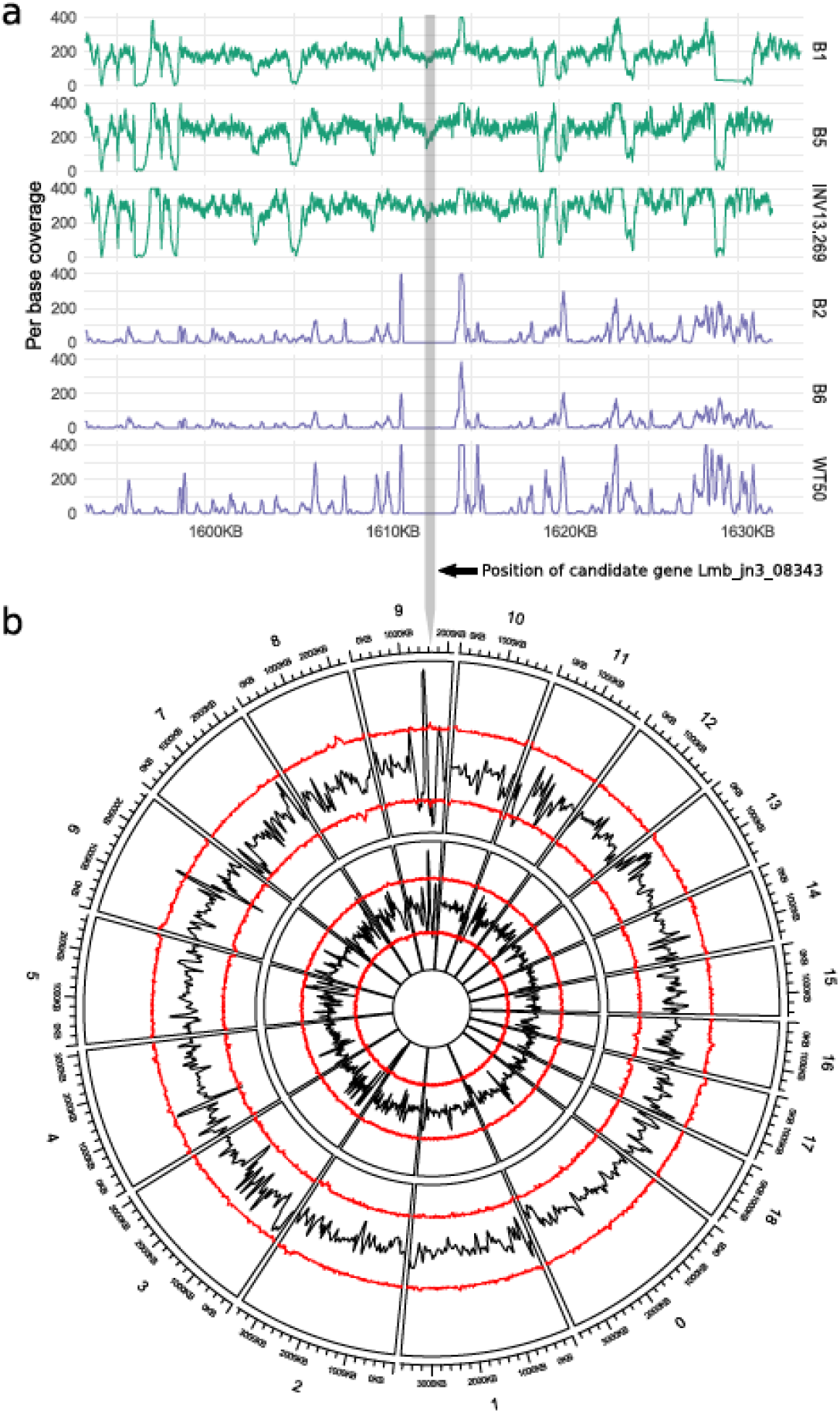
Identification of the candidate region for *AvrLmS* by bulked segregant sequencing. (a) Per base coverage for all samples (not including Bulk 3 and 4, *AvrLm7* and *avrLm7* respectively) for 20kb upstream and downstream of the gene Lmb_jn3_08343 (coding sequence demarcated with grey vertical bar) on scaffold 9 (JN3_SC09). Samples with *AvrLmS* are shown in green and those with *avrLmS* in purple. Y-axis limit was set to 400. (b) Circos plot of Δ(SNP-index) between Bulk 1/2 with *AvrLmS* (outer circle) and Bulk 5/6 with *AvrLmS*+*AvrLm7* (inner circle) for the 19 scaffolds larger than 1Mb. Confidence intervals of 99% (red) are shown. The y-axis is bounded from −0.5 to 0.8 for the outer plots and from −0.9 to 0.9 for the inner plots. The shared QTL for *AvrLmS* is located on scaffold 9. Coordinates for QTL are provided in Supplementary Tables S5 and S6.

**FIGURE 3.**
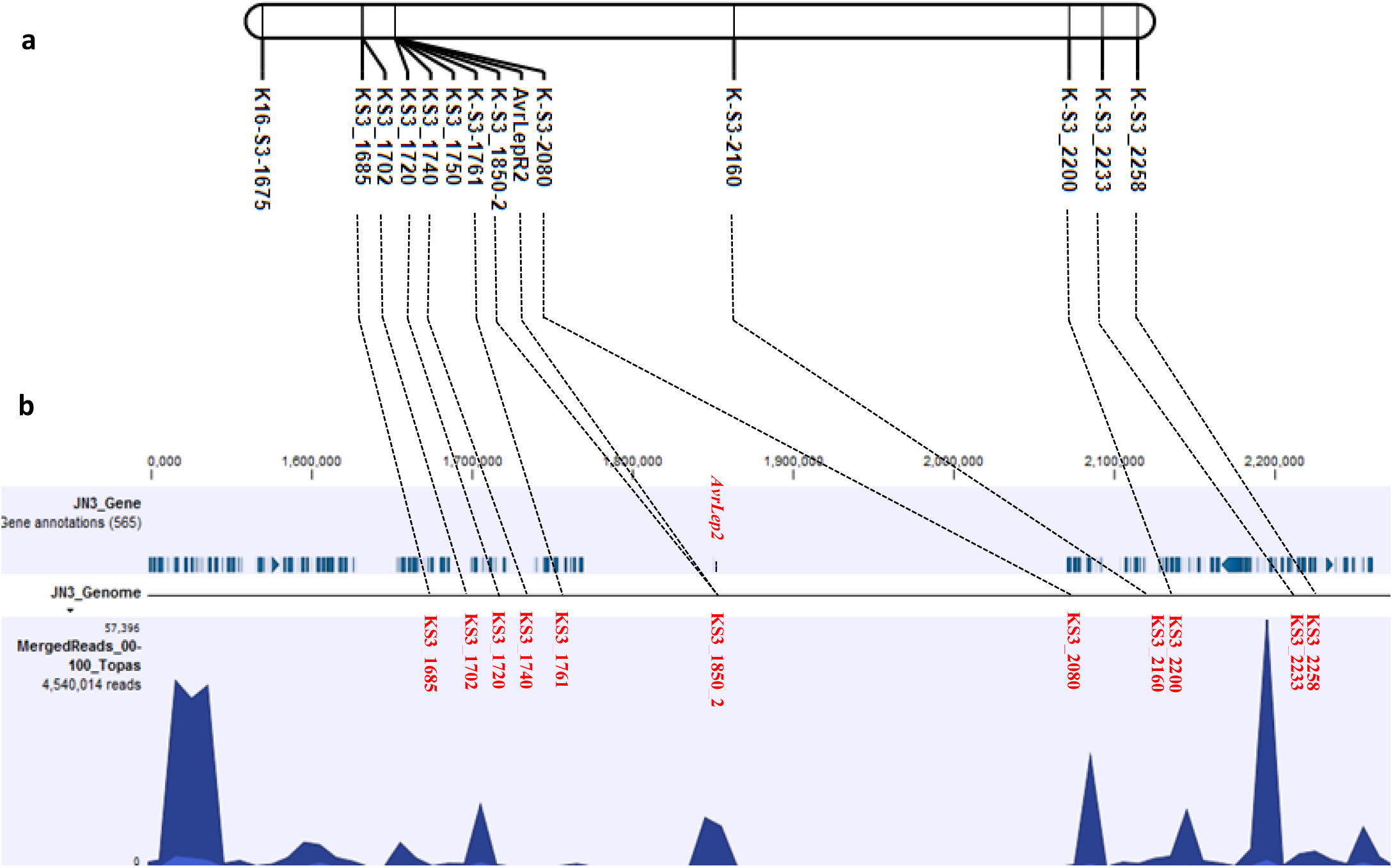
Genetic and physical maps of the *AvrLep2* genomic region in *Leptosphaeria maculans* isolate v23.1.3. (a) Position of *AvrLep2* relative to KASP markers on ‘00-100 x v23.1.3’ (SuperContig 3 from v23.1.3 V1 assembly GCF_000230375.1) map. (b) Physical region spanning the *AvrLep2* locus in the isolate v23.1.3. The top lane denotes predicted genes, bottom lane shows cumulative gene expression level for predicted *L. maculans* genes during infection time course (2-8 dpi).

#### Approach 2: Map-based identification of a candidate gene for *AvrLep2*

Under the conditions used for pathotyping at AAFC Saskatoon, the isolate v23.1.3 (JN3) produced an intermediate-virulent phenotype, with little to no plant response, on the cotyledons of plants carrying *LepR2* and was thus deemed ‘virulent’. The population of *L. maculans* F_1_ progeny produced from crossing v23.1.3 (*avrLep2*) and 00-100 (*AvrLep2*) segregated for virulence (41 isolates) and avirulence (57 isolates) towards *LepR2*, with the segregation ratio approximating a 1:1 ratio (*X*^2^ = 2.61, *P* = 0.11), as expected for genetic control of the phenotype by a single AVR gene. All 98 progenies were virulent on the susceptible line Topas DH16516. One hundred and fifty five KASP (Kompetitive Allele Specific PCR) markers were developed based on the whole genome sequence and predicted effector genes of *L. maculans* v23.1.3 and were applied to the progeny of the v23.1.3 × 00-100 cross. Two markers, K16-S3-1675 and K-S3-2160, closely segregated with the *AvrLep2* locus and spanned a physical interval of approximately 485 kb of the *L. maculans* genome. To more precisely map the *AvrLep2* locus, an additional eleven KASP markers were designed within the *AvrLep2* interval. The resulting map showed that *AvrLep2* resided within an interval of 319 kb between two markers, K-S3-1761 and K-S3-2080 (Figure 3). To improve the predicted gene annotation within the *AvrLep2* interval, previously generated RNA-Seq data produced from *L. maculans* infected *B. napus* seedling (Haddadi *et al*., 2016; 2019) were mapped to the *L. maculans* genome. Genes within the *AvrLep2* interval were manually annotated and a predicted secreted protein was identified as the AvrLep2 candidate.

#### Two approaches, one ‘typical’ avirulence effector gene candidate

The two cloning strategies identified the same candidate gene, Lmb_jn3_08343, which is 426 bp and contains one exon. In isolate v23.1.3, it is located in a typical AT-rich region of 285 Kb containing one single gene (Figure 3, Supplementary Figure 3). It encodes for a small (141 AA) putative secreted (SignalP 4.1, Petersen *et al*., 2011) protein enriched in cysteines (8 cysteine residues in the mature protein). PCR amplification confirmed the gene was absent in the virulent isolate WT50 and in all virulent isolates in X82 progeny, while sequencing of the gene in the avirulent isolate INV13.269 indicated it is 100% identical to that of v23.1.3.

We examined single nucleotide polymorphism (SNP) events within the candidate gene in the previously-resequenced genomes of 36 additional *L. maculans* isolates from the AAFC collection (5 *avrLep2* and 31 *AvrLep2* isolates; Ghanbarnia *et al*., 2015). The candidate gene was present in all isolates. In total, eight nucleotide changes were observed in the candidate gene of which four resulted in non-synonymous amino acid substitutions (Supplementary Table 8 and Supplementary Figure 4). Among the mutations only A^278^ was invariant in all avirulent isolates, while G^278^ was present in most virulent isolates.

**FIGURE 4.**
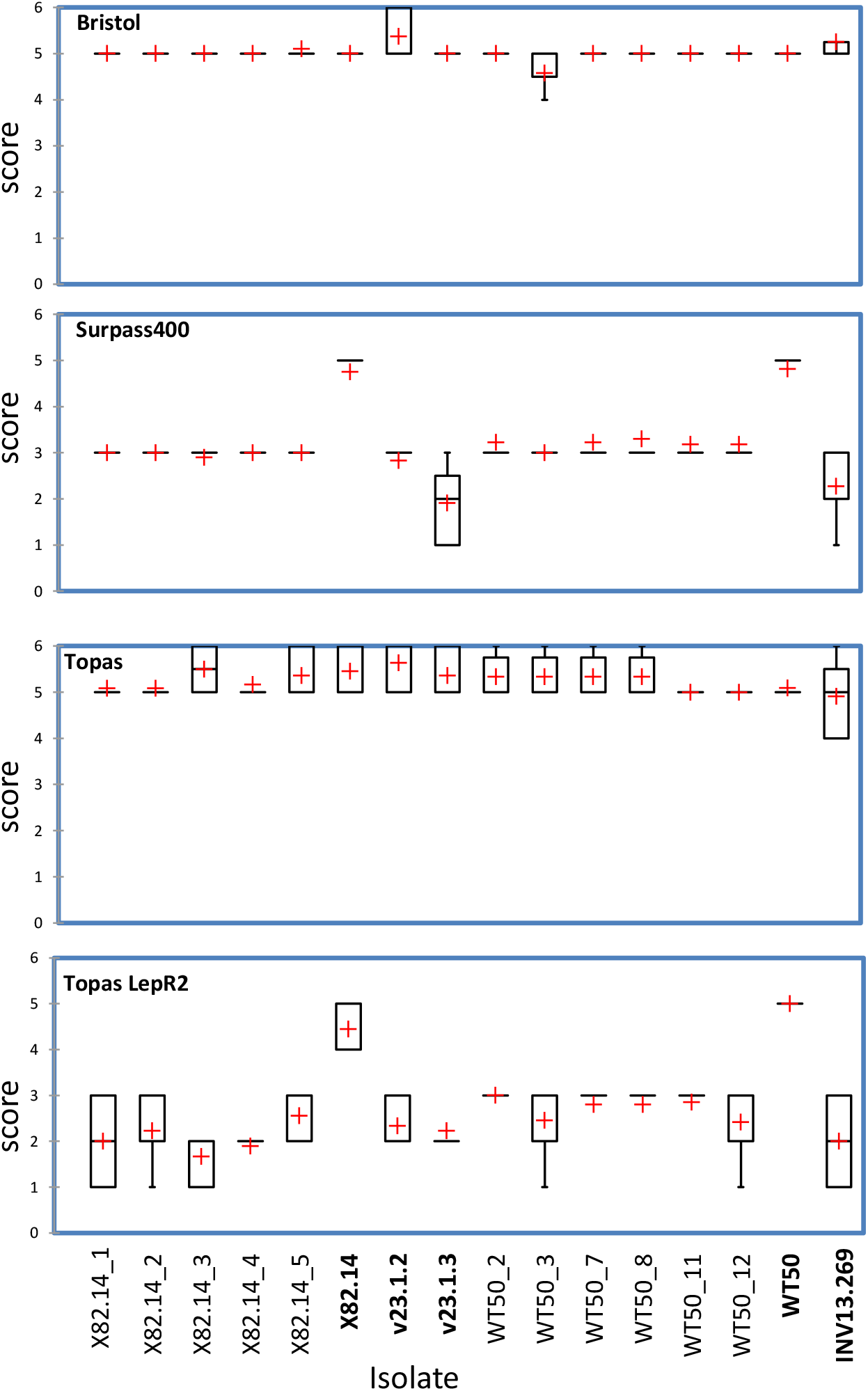
Box plot of rating scores of interaction phenotypes between wild-type or transformed isolates of *Leptosphaeria maculans* and four *Brassica napus* genotypes. From top to bottom: susceptible check Bristol (*Rlm2*-*Rlm9*), Surpass 400 (*LepR3-RlmS*), Topas DH16516 (no R gene), Topas-*LepR2* (*LepR2*). For each box, the red cross indicates the score mean; the black horizontal line, the score median, the rectangles comprise75 % (Q1-Q3) of the rating scores. Wild type isolates are in bold; X82.14-i and WT50-j are five and six independent transformed isolates of X82.14 and WT50, respectively, with the candidate gene Lmb_jn3_08343.

**FIGURE 5.**
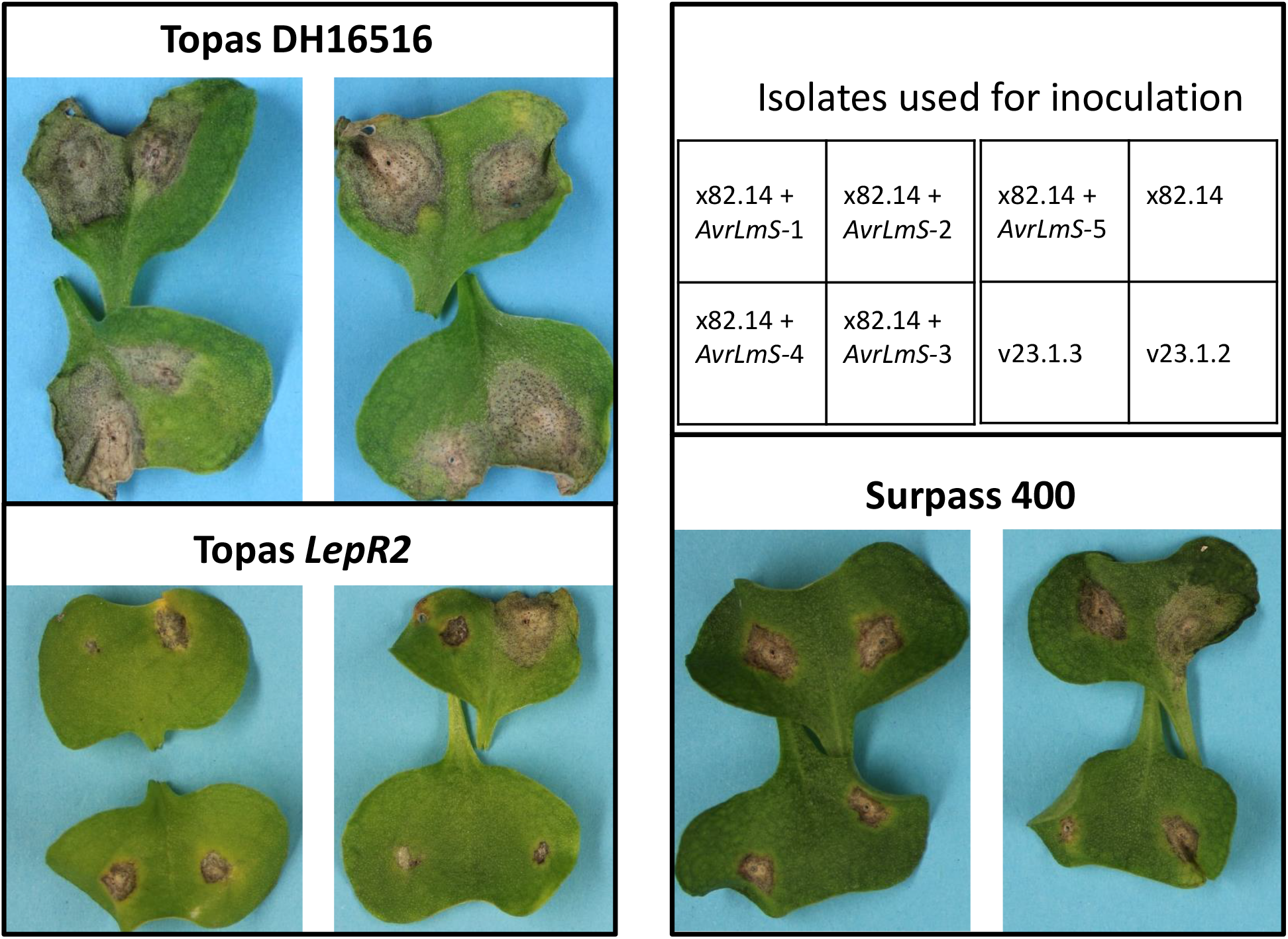
Examples of interaction phenotypes of wild-type and transformed isolates with the *AvrLmS* candidate gene on Topas DH16516, Topas-*LepR2* and Surpass 400. The four isolates inoculated on each plant are described on the up right panel. v23.1.3 and v23.1.2 are avirulent on Topas-LepR2 and Surpass 400 and X82.14 is virulent. All other isolates are independent transformants issued from the complementation of X82.14 isolate with the *AvrLmS* candidate gene. Pictures are taken 15 days post inoculation under BIOGER’s conditions.

### Validation of candidate genes

Two independent validation experiments were performed. First, WT50 (virulent on *RlmS* and *Rlm6*) and its progeny isolate X82.14 (virulent on *Rlm6, Rlm7* and *RlmS*), both deleted for the candidate gene, were complemented with the wild type copy (v23.1.3 allele) of the candidate gene (Lmb_jn3_08343). All complemented isolates remained fully virulent toward *Rlm1, Rlm2, Rlm3, Rlm4, Rlm6*, and *Rlm9* (Figure 4). In addition, X82.14 complemented isolates remained virulent towards *Rlm7*. All complemented isolates were found to induce the typical intermediate resistance of isolate INV13.269 on Surpass 400. Because of the presence of *AvrLm1* in v23.1.3, interacting with *LepR3* present in Surpass 400 (Larkan *et al*., 2013), the characteristics of the phenotype induced on Surpass 400 by the v23.1.3 Lmb_jn3_08343 allele could not be determined following inoculation with v23.1.3. However, the interaction phenotype of INV13.269 and of complemented isolates with the v23.1.3 allele confirmed this allele confers an intermediate phenotype on Surpass 400, which could develop toward susceptibility with time in some plants, as initially described (Van de Wouw *et al*., 2009). Finally, all complemented isolates were virulent on Topas DH16516 but displayed a clear resistant phenotype on Topas-*LepR2* (Figures 4-5). Therefore, Lmb_jn3_08343 encodes for the avirulence effector protein matching *RlmS* and is also able to elicit the *LepR2* resistance response.

Secondly, two types of construct for transformation using the *AvrLep2* candidate allele from *L. maculans* isolate 00-100 were produced, i.e. either with its native promotor, or with the promotor of the avirulence gene *AvrLm1*. After transforming the ‘virulent’ isolate v23.1.3 with the candidate gene constructs, restoration of avirulence phenotype was evaluated by inoculation of transgenic isolates on Topas*-LepR2* (Table 3). Transformant selections for each of the constructs were tested on the *B. napus* differential lines and showed avirulence on cotyledons of Topas-*LepR2* plants but remained virulent on the susceptible Topas DH16516 and Westar control lines (Figure 6). Positive transformants also showed wild-type interaction phenotypes with the differential lines harbouring other resistance genes (Table 3), confirming the identity and the specificity of the candidate gene as *AvrLep2*.

**FIGURE 6.**
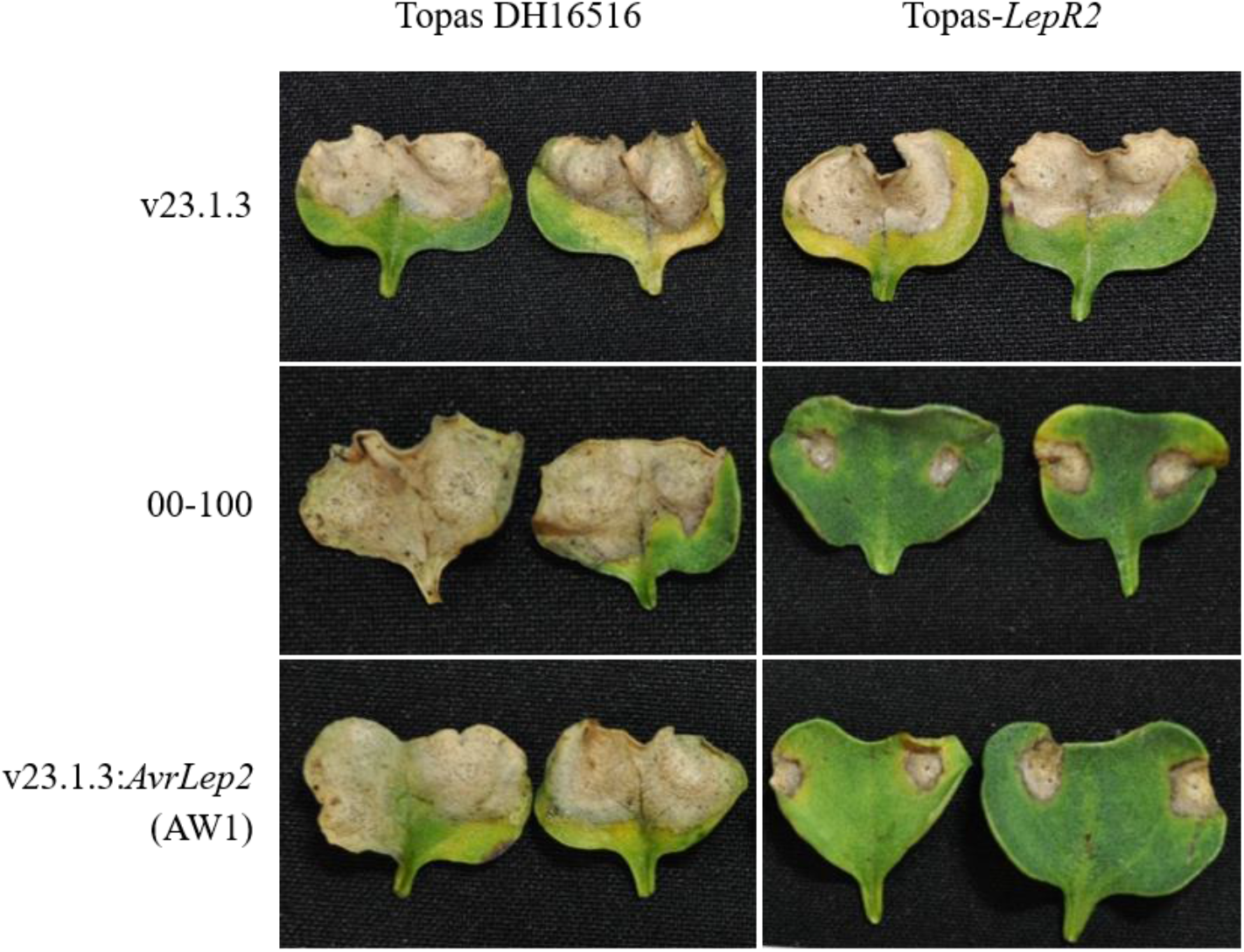
Phenotypic interaction of wild type and complemented *Leptosphaeria maculans* isolates on the cotyledons of control (Topas DH16516) and Topas-*LepR2* lines. Photographs of the infected cotyledons were taken 14 days post-inoculation. v23.1.3:*AvrLep2* (AW1) is a transformant with pNL11-*AvrLep2* construct (*AvrLep2* avirulent allele coding region was amplified from isolate 00-100, driven its native promoter).

### Expression Analysis

Previously generated RNA-Seq data with isolate v23.1.2, avirulent on Surpass 400 (Van de Wouw *et al*., 2009), were used to compare the expression kinetics of Lmb_jn3_08343 with that of all previously cloned *L. maculans* avirulence genes, following inoculation of cotyledons of a susceptible cultivar (Dutreux *et al*., 2018; Leontovyčová *et al*., 2020). Lmb_jn3_08343 is highly expressed during cotyledon infection, with a peak of expression seven days after infection (dai) in BIOGER’s controlled conditions, i.e. before symptoms develop. It is fully co-regulated with previously cloned avirulence genes, particularly with *AvrLm4-7, AvrLm5-9* and *AvrLm3* (Figure 7). Previously generated RNA-Seq data for the infection of the susceptible *B. napus* line Topas DH16516 by both v23.1.3 and 00-100 (Haddadi *et al*., 2015) was also examined to determine the expression patterns for both alleles of *AvrLep2*. Peak expression, measured as reads per kilobase of transcript per million mapped reads (RPKM), was observed at 4 dai for both v23.1.3 and 00-100, with *AvrLep2* having a similar expression to *AvrLm5-9* in both isolates (Supplementary Figure 5).

**FIGURE 7.**
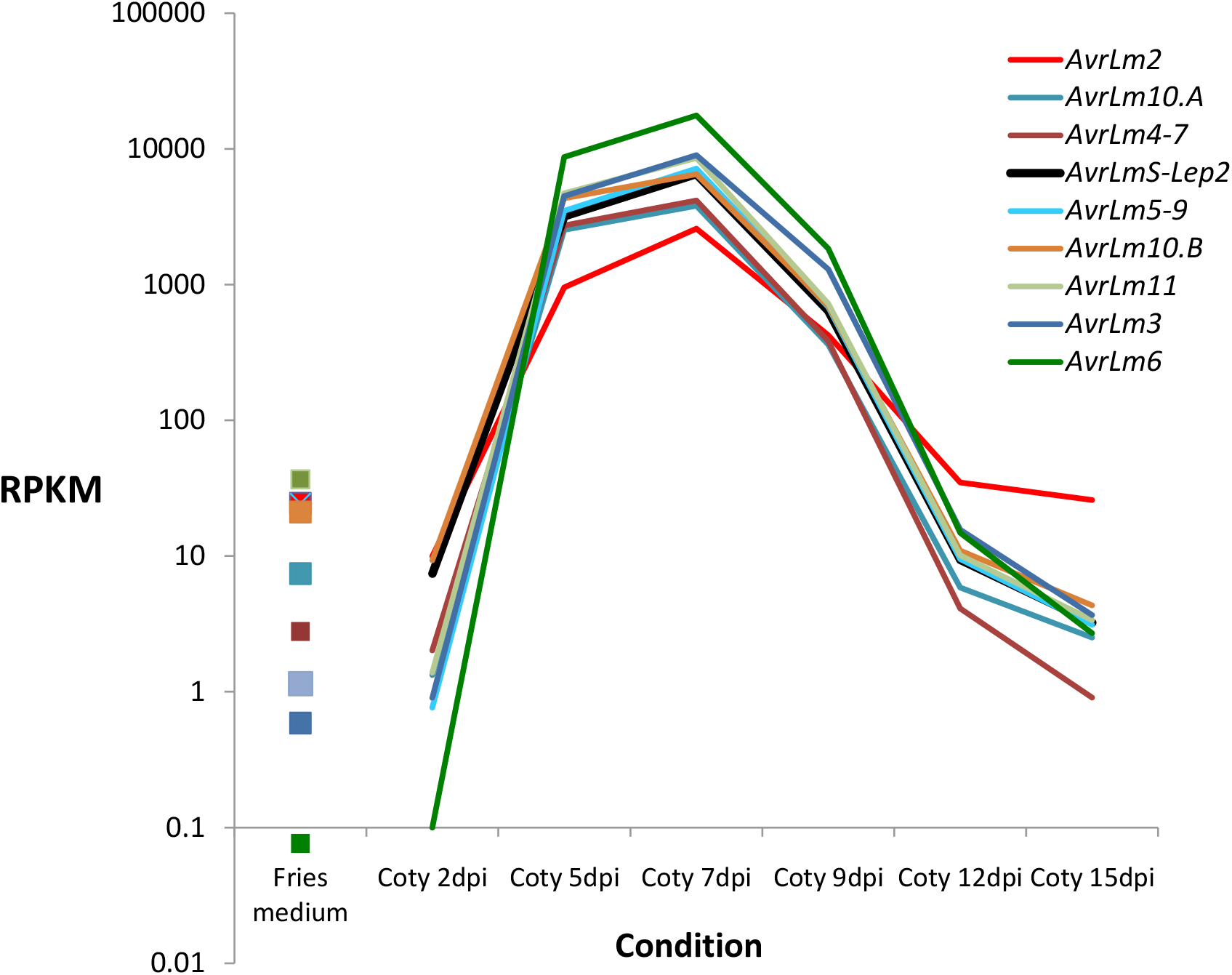
Expression of Lmb_jn3_08343 upon infection of oilseed rape cotyledons. Expression kinetics of Lmb_jn3_08343 was compared to that of previously cloned *AvrLm* genes. RNA-Seq data were obtained for isolate v23.1.2 *in vitro* (Fries medium condition) and following infection of cotyledons of cv. Darmor-*bzh* at 2, 5, 7, 8, 12 and 15 days post infection (dpi). Values are RPKM.

### Adult Plant Tests

After leaf infection and leaf spot development in the field, *L. maculans* grows systemically into the petioles and the stems before switching to necrotrophy and developing the stem canker symptom. To test the functionality of the *LepR2*-*AvrLep2* interaction during these later stages of plant infection, seedlings of Topas DH16516 and Topas-*LepR2* were inoculated with the control isolates v23.1.3 and 00-100, as well as eight additional native *L. maculans* isolates that had previously been classified as being ‘virulent’ towards both *LepR2* and *RlmS* based on cotyledon pathotyping. Three of the isolates had deletions of *AvrLep2*, while the remaining five all contained intact *AvrLep2* alleles of either the v23.1.3 (A^278^) or 00-100 (G^278^) haplotype, based on whole-genome resequencing (Supplementary Table 9). After allowing the infections to proceed from the cotyledon into the stem, where the lesion development was allowed to proceed for 12 weeks post-inoculation, there was a visible difference amongst the isolates in internal infection of Topas-*LepR2*, despite all of them (except the avirulent control 00-100) produced virulent cotyledon interactions. All seven isolates carrying an intact *AvrLep2* allele produced significantly less internal infection in the Topas*-LepR2* plants than in the susceptible Topas DH16516 control plants (Mann-Whitney test, *P* values ranging from 0.028 to <0.0001) (Figure 8, Supplementary Table 9). Only one ‘*AvrLep2*’ isolate (AI397) was able to produce relatively high infection in Topas-*LepR2*. Further analysis of the *AvrLep2* allele for this isolate revealed a novel substitution (C^56^ -> T^56^) unique amongst all sequenced isolates. In contrast, all three isolates which carried a deletion at the *AvrLep2* locus (B16-13, B18-10 and B18-11) produced high and identical levels of infection in both the control Topas DH16516 and Topas-*LepR2* lines.

**FIGURE 8.**
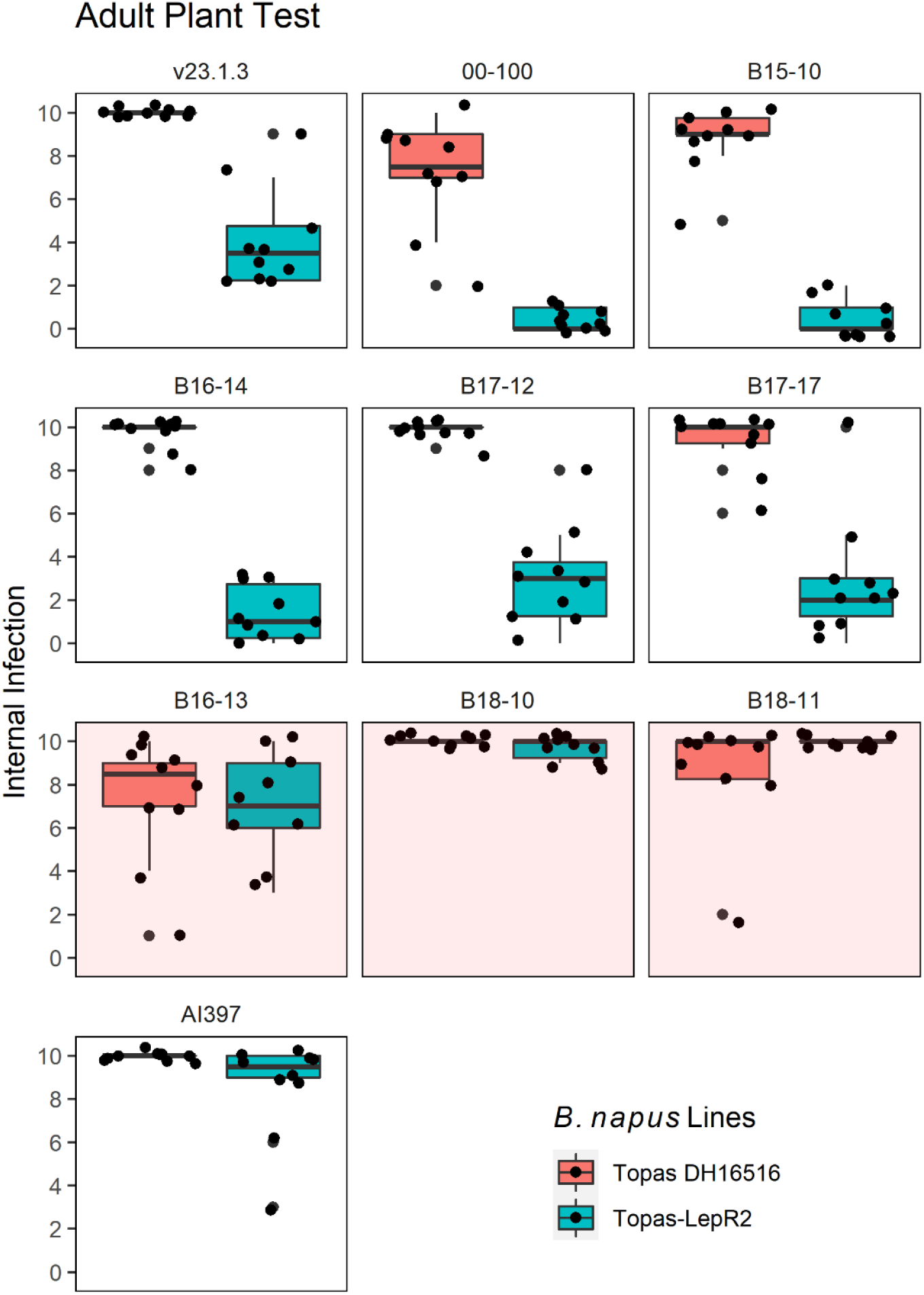
Box & Whisker Plot for internal infection of Topas DH16516 (red) and Topas-*LepR2* (blue) of adult plants by 10 *L. maculans* isolates. Data points (black dots) indicate internal infection (0-10 scale) of individual plants (10 per test). Boxes denote interquartile range (Q_1_ to Q_3_), black bars show median score and whiskers denote range of distribution. Data points outside the whiskers (< 1.5 Q3 or >1.5 Q1) are considered outliers. Red-shaded panels indicate isolates for which *AvrLep2* is deleted.

### A posteriori control of the consistency of phenotypic data

The sequenced reference isolate v23.1.3 had been shared between laboratories but here, interaction phenotypes of v23.1.3 inoculated on Topas-*LepR2* clearly differed between AAFC and BIOGER experiments in spite of the use of the same Topas-*LepR2* seed lot. To resolve this difference, the two v23.1.3 lines maintained for years in parallel at AAFC and BIOGER were shared again and phenotyped on differential plant genotypes including Topas-*LepR2* at BIOGER. The interaction phenotypes of the two clonal isolates on Topas-*LepR2* were identical, with a clear resistance response compared to virulent control isolates (Supplementary Figure 6). In addition, the two isolates behaved similarly on all other plant genotypes including those containing resistance genes *Rlm1* or *Rlm4*, matching *AvrLm1* and *AvrLm4-7* present in v23.1.3. This suggests that only environmental conditions or experimental parameters, not genetic drift after independent subculturing of the isolate in the two laboratories, explain the difference in the phenotypic expression of the *LepR2 /AvrLmS-AvrLep2* interaction.

## DISCUSSION

In search of the avirulence genes matching *RlmS* in Surpass 400 and *LepR2* in Topas-*LepR2*, (i.e., *AvrLmS* and *AvrLep2*, respectively) we report here on the independent cloning by two teams of the same avirulence gene, in spite of clear divergent interpretations of the interaction phenotypes observed on resistant plant genotypes between the two laboratories. Cloning *AvrLmS* is the first example of applying the BSS strategy to clone a gene of interest from *L. maculans*. BSS is a powerful approach to rapidly identify candidate genes not only in plant species (Klein *et al*., 2018, Dong *et al*., 2018) but also in fungi (Lenhart *et al*., 2019; Hu *et al*., 2015). Here, we validated BSS on the previously cloned *AvrLm4-7* gene and our cloning of *AvrLmS* indicates that small-sized bulks containing only ∼10 isolates are sufficient to identify the genomic region containing the candidate gene. On the other hand *AvrLep2* was cloned using the conventional bi-parental mapping approach. Similar to all other *AvrLm* genes, *AvrLmS*-*AvrLep2* is located in an AT-rich genome environment, encodes for a small secreted protein rich in cysteines, and the gene is highly overexpressed at early stages of cotyledon infections. The gene was renamed here *AvrLmS-Lep2*.

The interaction phenotype of the reference isolate v23.1.3 on Topas-*LepR2* was interpreted as either an intermediate resistance phenotype, or a ‘virulence’ phenotype. Such intermediate phenotypes have been reported for this interaction in many studies. First, the phenotypic resistance response on the *B. napus* cultivar Surpass 400 inoculated with avirulent *AvrLmS* isolates was described as intermediate, i.e. producing larger lesions than typical hypersensitive response (HR), and sometimes at the edge of virulence, depending on the environmental conditions or with time (Van de Wouw *et al*., 2009). On the plant side, the resistance in Surpass 400 was at first considered as monogenic, based on field assays in Australia (Li and Cowling, 2003) or genetic mapping (Yu et *al*. 2008), though later mapping with defined isolates under controlled conditions suggested the occurrence of two genes in Surpass 400 (named as *BLMR1* and *BLMR2*, Long *et al*., 2011; or *LepR3* and *RlmS*, Larkan *et al*., 2013). The resistance gene *LepR2* in DH line AD49 was described as limiting, but not preventing, hyphal growth of avirulent isolates, along with restricting sporulation on the infected plant tissues (Yu *et al*., 2005). In addition, following inoculation of a range of isolates, most of them (22 out of 32) were found to display large, non-sporulating lesions (scores between 3 and 6 on a 0-9 scale), while 8 isolates only displayed typical HR (score <3) (Yu *et al*., 2005). In the study by Yu *et al*. (2005) the two most virulent isolates on *LepR2* displayed scores between 6 and 6.5, but never reached the level of virulence observed on the susceptible genotypes (scores >7.5). This intermediate resistance phenotype was nevertheless correlated with the stem canker resistance (Yu *et al*., 2005), suggesting it is sufficient to prevent *L. maculans* systemic growth in the leaves and stems. Similarly, *B. napus* plants harbouring the resistant gene *BLMR2* (*LepR2*-*RlmS*) derived from the Surpass 400 parent also showed intermediate resistance response at the cotyledon stage (Long *et al*., 2011), and correlated with partial resistance response at the adult plant stage (Dandena *et al*., 2019), while *BLMR1*, corresponding to *LepR3* (Dandena *et al*., 2019), gave a strong and typical HR.

Consistent with these published data, such an intermediate resistant phenotype on Surpass 400 was also described here for either the avirulent parental isolate INV13.269 or the transgenic isolates complemented with the v23.1.3 allele of *AvrLmS*). The fluctuating intermediate phenotype resulting from *AvrLmS-RlmS* or *AvrLep2*-*LepR2* interaction could be attributed to sequence variation in the gene, with only deleted *AvrLmS-Lep2* alleles, like those found in isolate WT50 or all virulent progeny of cross #82, able to induce clear susceptibility symptoms on Surpass 400 or Topas-*LepR2*, while variations in nucleotide sequence of the gene could correspond to variable degrees of avirulence, ranging from strong resistance for isolate 00-100 to intermediate resistance (or ‘virulence’) in v23.1.3. However, no clear relationship between sequence variants and phenotype was observed amongst sequenced isolates. Alternatively, the variation in cotyledon phenotypes observed in AAFC tests may be due to the expression level of the gene in v23.1.3 and its progeny. *AvrLep2* is expressed at a lower level than some other *AVR*, and at a similar level to *AvrLm5-9*, another *AVR* gene which generally also elicits an intermediate resistance response (Ghanbarnia *et al*., 2015) that can be challenging to identify through cotyledon phenotyping. Another hypothesis to explain the variable expression of symptoms and contrasting interpretation of the interaction could be a strong influence of environmental conditions on the phenotypic outcome, resulting in an ‘intermediate virulence’ phenotype in AAFC environmental conditions. Previous studies have shown the impact of temperature or humidity on the expression of some *AvrLm*-*Rlm* interactions (for example, Huang *et al*., 2006). Consistent with that, the avirulent phenotype of v23.1.3 on Topas-*LepR2* observed at BIOGER was reproduced here under BIOGER’s conditions using seed lots and isolate used at AAFC. Importantly, regardless of the cotyledon phenotype, it was clearly shown that the presence of an *AvrLep2* allele in any isolate was sufficient to induce *LepR2* dependant resistance in the adult plant assay, with variable but significant reduction of stem necrosis, while a deletion of the *AvrLep2* gene always resulted in similar internal infection and stem lesion in the *LepR2* line as in the susceptible control (Figure 8).

In spite of divergent interpretation of the phenotypes, the use of crosses involving isolates with contrasting phenotypes on the resistant plant genotype allowed us to identify and validate Lmb_jn3_08343 as the matching avirulence gene. Both approaches involved crosses between isolates displaying differential phenotypes on Surpass 400 and/or Topas-*LepR2*, with either a highly susceptible X intermediate resistance combination (WT50 x INV13.269), or an intermediate ‘virulence’ X avirulence combination (v23.1.3 x 00-100) that finally targeted the same avirulence gene. Understanding the relationship between allelic variation and interaction phenotypes, and how environmental or experimental conditions, along with the effect of the plant genetic background, can modulate the outcome of the interaction should be further analysed in future work. Supported by our phenotypic data, we thus showed that the genes identified in Surpass 400 as *RlmS, LepR2* or *BLMR2* recognize the same effector protein and are likely the same resistance gene. The name *LepR2*, initially published in the literature, should be retained for the resistance gene, while the corresponding effector gene has now been renamed *AvrLmS-Lep2*. The different nomenclatures defining the *R* genes in Surpass 400 in the past have now been clarified through this study, with LepR3 interacting with AvrLm1 and LepR2 interacting with AvrLmS-Lep2. This work illustrates a first step toward the standardization of the complex and divergent terminologies used to describe *L. maculans* – *Brassica* sp. interactions.

## EXPERIMENTAL PROCEDURES

### Approach 1: Bulk Segregant Analysis

#### *L. maculans* isolates and crosses

To map *AvrLmS*, a segregating progeny population was built following an *in vitro* cross between isolate WT50, isolated in Australia in 2005 (van de Wouw *et al*., 2009), and INV13.269, recovered in 2013 in France. *In vitro* crosses and random ascospore progeny recovery were performed as previously established (Plissonneau *et al*., 2016).

#### Plant genotypes and inoculation tests

Isolates were grown on 20% V8-agar medium to produce conidia. Conidia (10 µL, 10^7^ spores mL^-1^) were inoculated onto wounded cotyledons of 10 to 12 10-days old seedlings per plant genotype. The following *B. napus* plant genotypes were used: Westar or Topas DH161516 (no *R* gene), 15-23-4-1 (*Rlm7*), Pixel (*Rlm4*), Columbus (*Rlm1, Rlm3*), Darmor (*Rlm9*), Bristol (*Rlm2*-*Rlm9*), Darmor-MX (*Rlm6*-*Rlm9*), 15.22.5.1 (*Rlm3*) (Balesdent *et al*., 2005), Topas-*LepR2* (*LepR2*; Larkan *et al*., 2016) and Surpass 400 (*LepR3-RlmS*; Larkan *et al*., 2013). Four different isolates were inoculated on each plant. Symptoms were scored two to three times between 12-21 dai, using a 1-6 scale, with scores 1-3 and 4-6 corresponding to avirulent and virulent phenotypes, respectively (Balesdent *et al*., 2005). To compare the interaction phenotypes of progeny isolates with those of parental isolates, the non parametric Kruskall-Wallis test was applied, with a *P* value threshold set up at 0.05, using XLSTAT Version 2013.4.03. The phenotypes of the progeny selected for BSS were confirmed in an independent inoculation test.

#### DNA extraction and bulk preparation

Isolates were grown on Fries liquid medium for seven days as previously established (Fudal *et al*., 2008). Mycelium was harvested by vacuum filtration, rinsed with sterile de-ionized water and freeze-dried. DNA was then extracted using the DNeasy Plant Mini Kit (Qiagen,Hilden, Germany) following manufacturer’s instructions. DNA concentration was quantified using Qubit™ dsDNA BR Assay Kit (Invitrogen, Carlsbad, USA). For library preparation, each isolate sample was adjusted to 20 ng of DNA and pooled into six different bulks (Table 2). The final DNA concentration in the bulks was controlled similarly.

#### Whole genome-sequencing

Between 74 and 200 ng of DNA was taken from each sample to prepare the DNA library. The DNA library was prepared using the Illumina Nextera™ DNA Flex Library Prep Kit (Illumina Inc., San Diego, USA) according to the manufacturer’s protocol. Whole genome sequencing was performed on each of the eight bulks using the Illumina Hi-Seq technology with 150 bp PE at Kinghorn Centre for Clinical Genomics (KCCG) Core Facility at the Garvan Institute of Medical Research (Darlinghurst, Australia).

#### Read mapping and variant calling

Quality trimming of reads was carried out using Trimmomatic 0.36 (Bolger *et al*., 2014) with default parameters and the Nextera paired-end adapters provided with the software. Reads were mapped to the reference isolate v23.1.3 (GenBank BioProject: PRJEB24468, Assembly GCA_900538235), using BWA 0.7.17 with the BWA-MEM algorithm (Li, 2013) and default parameters. Duplicates were removed using the Picard MarkDuplicates 2.8.1 (Picard Toolkit). Reads with mapping quality <20 were filtered using SAMtools 1.8 (Li *et al*., 2009). Variants were called using GATK HaplotypeCaller v3.6-0-g89b7209 (McKenna *et al*., 2010) with default parameters. Paired bulked samples were extracted (Bulk 1/2, Bulk 3/4, Bulk 5/6) using VCFtools 0.1.15 (Danecek *et al*., 2011) and variants with a phred-scaled quality score <30 were excluded. Indels were removed using GATK SelectVariants. The public reference genome repeat annotation was used to exclude SNPs occurring within repeats. SNPs that were heterozygous or monomorphic in parental isolates were also excluded. Finally, VCF files were converted to tabular format with GATK VariantsToTable.

#### QTL-seq and candidate SNP analysis

QTL-seq of paired bulked segregants was carried out using the R package QTLseqr version 0.7.3 (Mansfeld and Grumet 2018; Takagi *et al*., 2013) with the QTL-seq approach (Takagi *et al*., 2013). For SNP filtering settings, we used minTotalDepth = 100, maxTotalDepth = 800 and a minimum genotype quality of 99. Reference allele frequency was required to be >=0.2 and <= 0.8. Window size was set to 5e^4^. To complement this analysis, SNPs that segregate perfectly between *AvrLmS* (Bulk 1 and 2 including the parent INV13.269) and *avrLmS* (Bulk 5 and 6 including the parent WT50) were identified as candidate SNPs. We consider a SNP as perfectly segregating if it is called as homozygous by GATK in all samples and the alleles differ between samples with *AvrLmS* and those with *avrLmS*. The candidate SNP positions were intersected with the gene annotation to identify candidate genes based on the presence of a candidate SNP in the gene sequence or in the 5kb upstream/downstream region. The candidate region was also queried for long terminal repeats (LTR) using RepeatMasker. The GC content in the candidate QTL region was analysed using seqinr 3.4 (Charif and Lobry 2007) and AT-rich regions were identified with OcculterCut 1.1 (Testa et al., 2016).

#### Candidate gene analysis

Gene presence/absence variation (PAV) analysis was performed on the *AvrLmS* candidate gene. To assess PAV, the SAMtools view utility was used (Li *et al*., 2009). Per base coverage of the candidate gene and upstream and downstream regions was calculated using BEDTools 2.26.0 (Quinlan and Hall 2010) and plotted with ggplot2 in R. To search for gene homology of the *AvrLmS* candidate gene, the full nucleotide sequence of the candidate gene was BLAST queried against the InterProScan database website. The genomic region surrounding the candidate gene including 10 kb upstream and downstream regions was also queried using BLAST on the NCBI database website. Expression of *AvrLmS* was examined from infection time course data previously generated following inoculation of isolate v23.1.3 on the susceptible cv. Darmor-*bzh*, or from *in vitro* culture conditions (Dutreux *et al*., 2018; Leontovyčová *et al*., 2020).

#### Functional validation of the candidate gene

The *AvrLmS* candidate gene was amplified from genomic DNA of v23.1.3 (2537-bp fragment: 1049-pb upstream and 1062-bp downstream of the CDS) and cloned into the binary vector pPZPNat1 using Gibson assembly (New England Biolabs, Ipswich, USA). Plasmid was amplified in *Escherichia coli* TOP10 cells, re-extracted and checked by sequencing (Eurofins Genomics, Ebersberg, Germany). The construct was introduced into *Agrobacterium tumefacien*s strain C58 by electroporation at 1.5 kV, 200 ohms and 25 lF and used for transformation of two virulent isolates, WT50 and X82.14, as described by Gout *et al*. (2006). Fungal transformants were selected on 50 µg mL^-1^ nourseothricin (WERNER BioAgents, Jena, Germany), purified by single pycnidium isolation and maintained on selective medium. 21 and 7 independent transformants were recovered for WT50 and X82.14, respectively. To control the deletion of the candidate gene in WT50 and in virulent progeny, the primers AvrLms-up (GACTGCAACACCTCTTTTCCA) and AvrLms-low (CGCTCGATCCGTCCCTTATA) were used on genomic DNA using standard PCR procedures and an annealing temperature of 60°C.

### Approach 2: Map-based cloning

#### Phenotyping of mapping population

For mapping *AvrLep2*, a F_1_ population produced from the parental isolates v23.1.3 and 00-100, previously used to map the *AvrLm5-9* locus (Ghanbarnia *et al*., 2018) was also shown to be segregating for the *AvrLep2* phenotype under the controlled growth chamber conditions used at AAFC Saskatoon (Larkan *et al*., 2013). The *B. napus* line Topas-*LepR2* (Larkan *et al*., 2016) and the *LepR2* line 1135 (Yu *et al*., 2009) were used to determine the phenotypic response of the parental isolates and progeny to *LepR2. B. napus* cotyledons were inoculated as described previously (Chen and Fernando 2006). Each *L. maculans* isolate was tested on 12 seedlings of the differential lines and 12 seedlings of Topas as susceptible control. The disease reactions were scored 14 dai and rated using the 0-9 scale described by Williams (1985). Paired-end Illumina sequencing and assembly of parental isolates was previously described by Ghanbarnia *et al*. (2015).

#### Expression analysis by RNA-Seq

Expression of *AvrLep2* was examined from infection time course data previously generated (Haddadi et al., 2016). Briefly, cotyledons of 7-day-old Topas DH16516 seedlings were inoculated with the parental isolates 00-100 and v23.1.3. Mock inoculation with water served as a negative control. Cotyledon discs 6 mm in diameter were excised from the infected cotyledons (four biological replicates) at 2, 4, 6 and 8 dai. RNA was extracted and sequence reads (100 bp paired-end) were generated with Illumina TruSeq high output version 3 chemistry on a HiSeq 2500 (Illumina, Inc.) at NRC-Plant Biotechnology Institute (NRC*-*PBI), Saskatoon, Canada.

#### Mapping, cloning and transformation of candidate gene

SNPs for primer development were selected based on whole genomic comparison of parental isolates or based on predicted polymorphic effectors from isolate 00-100 and v23.1.3 using CLC Genomic Workbench (version 8.1.1, CLC Bio; Denmark). Then the target SNP(s) were used to design the KASP primers using the PrimerPicker software provided by KBioscience (http://www.kbioscience.co.uk/). KASP reactions were performed as per the manufacturer’s instructions (LGC Biosearch; https://www.biosearchtech.com). One hundred F_1_ progeny were selected in order to screen KASP markers spanning the whole *L. maculans* genome (Rouxel *et al*., 2011). A linkage map of *AvrLep2* was constructed using MAP function of QTL IciMapping v3.2 software (Li *et al*., 2008). Minimum LOD (logs of the odds ratios of linkage vs. no linkage) scores of 6.0 (maximum recombination fraction of 0.6) were used to group loci. After initial linkage between markers and the *AvrLep2* locus was established, additional KASP markers targeted to the *AvrLep2* interval were designed based on genomic polymorphisms to enrich the map. Cloning, transformation and functional validation of *AvrLep2* SNP variants was performed as described previously (Ghanbarnia *et al*., 2015). For functional validation two constructions were produced. First, the ORF for *AvrLep2* candidate gene (426 bp) was amplified and transferred to the fungal transformation vector pLM4 (Ghanbarnia *et al*., 2015) under the control of the *AvrLm1* promoter. In addition, an *AvrLep2* candidate gene amplicon, including the native promoter region (starting from 1996 bp upstream of the ATG start codon based on the v23.1.3 reference sequence) and 186 bp downstream of the predicted ORF (total length 2609 bp), from the *AvrLep2* parental isolate 00-100 was transferred into the fungal transformation vector pNL11 (Larkan *et al*., 2013). To confirm the *AvrLep2* specificity, phenotypic response of the parental isolates and positive transformants (showing restored phenotypic reaction on Topas-*LepR2*) were tested using the following *B. napus* differential lines; Topas-*Rlm1*, Topas-*Rlm2*, Topas-*Rlm4*, Topas-*LepR1*, Quantum (*Rlm3*), Roxet (*Rlm7*), Goéland (*Rlm9*) and the *B. juncea* line Vulcan-1S (*Rlm6*) (Larkan *et al*., 2016). Topas DH16516 (no R genes) was used as a positive control for infection by *L. maculans*.

#### Adult plant tests

Isolates carrying different alleles of *AvrLep2* (v23.1.3-type, 00-100-type or deletion, as well as one unique mutation) and previously classified as ‘virulent’ towards both *LepR2* (Topas-*LepR2* line) and *RlmS* (72-1; a F3 progeny selected form the Topas DH16516 x Surpass 400 population (Larkan et al., 2013) which retains *RlmS* resistance but lacks *LepR3*) were used to infect Topas DH16516 and Topas-*LepR2* seedlings via standard cotyledon wounding method. The plants were maintained under controlled conditions (Haddadi *et al*., 2019) and infection was allowed to progress into the stem (cotyledons were not removed). The resistance phenotype was scored in the adult plants via assessment of internal infection in the stem at 8-12 weeks post-infection. Stem infection was rated using a 0-10 scale, where each graduation corresponds to 10% of the internal cross-section showing infection damage. Results were plotted using the ggplot2 (Wickham, 2016) and reshape2 (Wickham, 2007) packages in R 4.0.0 (R Core Team, 2020), run in RStudio v1.3.959.

## ACKNOWLEDGMENTS

This research was funded by the French National Research Agency project AvirLep (ANR GPLA07-024C) and the Grains Research and Development Corporation. The Canadian researchers acknowledge the generous funding received through the Growing Forward 2 Research Program and SaskCanola. We thank Régine Delourme, Pascal Glory (INRAE, UMR IGEPP) and the CRB BRACYSOL for providing the Darmor-MX line and Ralph Lange (Alberta Innovates) for providing *L. maculans* isolate AI397.

## AUTHOR’S CONTRIBUTION

TXN, KG, NJL, BO and ASE conducted the experiments; AS and PH performed bioinformatics analysis; TXN, AS, BO, KG, MHBo, NJL, MHBa and TR analysed the data; JB, TR, KG, NJL, MHBa, MHBo and TXN conceived the idea; JB, TR, and MHBa supervised the *AvrLmS* project, MHBo and WGDF supervised *AvrLep2* project; MHba coordinated the writing of the publication.

## SUPPORTING INFORMATION LEGENDS

**Supplementary Table 1**. Read mapping statistics obtained from library preparation of six individual bulk samples (samples 1-6) and parental isolates (samples 7 and 8).

**Supplementary Table 2**. Single nucleotide polymorphisms (SNPs) found in six bulked progeny. SNPs that were monomorphic, not homozygous in both parents, or that occurred in repetitive regions were excluded.

**Supplementary Table 3**. QTL-Seq analysis results for Bulk 3/4, showing QTL peaks passing the 99% confidence threshold and supported by >3 SNPs.

**Supplementary Table 4**. Merged QTL-Seq analysis results for all bulked progeny. Neighboring QTL within 30kb were merged.

**Supplementary Table 5**. QTL-Seq analysis results for Bulk 1/2, showing QTL peaks passing the 99% confidence threshold and supported by >3 SNPs.

**Supplementary Table 6**. QTL-Seq analysis results for Bulk 5/6, showing QTL peaks passing the 99% confidence threshold and supported by >3 SNPs.

**Supplementary Table 7**. AT-rich (R0) and GC-equilibrated (R1) regions on JN3 scaffold 9, identified by OcculterCut.

**Supplementary Table 8**. Allelic variation at the *AvrLep2* locus in a collection of 37 isolates virulent or avirulent towards *LepR2* and polymorphic sites in its protein.

**Supplementary Table 9**. Cotyledon and adult plant inoculation tests with virulent and avirulent isolates toward *LepR2*.

**Supplementary Figure 1**. Plot of Δ(SNP-index) between Bulk 3 and Bulk 4 (*AvrLm7* and *avrLm7*) across scaffold JN3_SC03. Confidence intervals of 95% (blue) and 99% (red) are shown. QTL coordinates are provided in Supplementary Table 3.

**Supplementary Figure 2**. Plot of GC content calculated in 500 bp windows across scaffold 9. The grey region demarcates the candidate regions identified using QTL-Seq for Bulk 1/2 and Bulk 5/6. The Red line showes the CDS of the candidate gene Lmb_jn3_08343.

**Supplementary Figure 3**. Genomic location and characteristics of the *AvrLmS* (Lmb_jn3_08343) gene in *Leptosphaeria maculans*

**Supplementary Figure 4**. Nucleotide sequence of the 486 nucleotide region encoding *AvrLep2* from *L*. maculans isolate 00-100 and its predicted amino acid sequence.

**Supplementary Figure 5**. Comparison of *Avr* gene expression in *L. maculans* isolates v23.1.3 (A) and 00-100 (B) during cotyledon infection of *B. napus* line Topas DH16156.

**Supplementary Figure 6**. Comparison of the pathogenicity of the two batches of v23.1.3 (JN3) from AAFC and from BIOGER.

