## SUPPORTING INFORMATION for "Two independent approaches converge to the cloning of a new *Leptosphaeria maculans* avirulence effector gene, *AvrLmS-Lep2*"

**Supplementary Table 1.** Read mapping statistics obtained from library preparation of six individual bulk samples (samples 1-6) and parental isolates (samples 7 and 8).

| Sample No. | Sample name | No. of isolates | No. of raw reads | No. of trimmed reads | Coverage depth (X) | Non-zero coverage depth[a] (X) | Horizontal genome coverage[b] (%) |
|---|---|---|---|---|---|---|---|
| 1 | AS | 25 | 47,134,448 | 46,981,574 | 164.56 | 173.00 | 95.12 |
| 2 | aS | 22 | 69,460,390 | 69,230,358 | 245.15 | 257.36 | 95.25 |
| 3 | A7 | 24 | 75,776,452 | 75,546,277 | 272.20 | 285.60 | 95.31 |
| 4 | a7 | 23 | 59,091,833 | 58,884,100 | 200.36 | 210.37 | 95.24 |
| 5 | AS7 | 11 | 66,939,344 | 66,711,323 | 215.49 | 226.04 | 95.33 |
| 6 | aS7 | 9 | 50,241,166 | 50,023,991 | 165.14 | 173.86 | 94.99 |
| 7 | INV13.269 (AS) | 1 | 65,658,080 | 65,475,808 | 242.12 | 256.58 | 94.36 |
| 8 | WT50 (aS) | 1 | 56,457,493 | 56,254,313 | 205.73 | 218.66 | 94.09 |

[a] Average number of reads per genome base, excluding bases with no reads aligned

[b] Number of bases in the genome covered by at least one read divided by total number of bases

**Supplementary Table 2.** Single nucleotide polymorphisms (SNPs) found in six bulked progeny. SNPs that were monomorphic, not homozygous in both parents, or that occurred in repetitive regions were excluded.

| | **Genotype call** | | | | |
|---|---|---|---|---|---|
| **Sample No.** | **INV13.269 allele** | **WT50 allele** | **Heterozygous allele** | **Missing** | **Total SNPs** |
| 1 | 9,123 | 10,768 | 44,414 | 1,422 | 64,305 |
| 2 | 9,762 | 9,502 | 45,407 | 1,056 | 64,671 |
| 3 | 9,353 | 9,787 | 45,625 | 962 | 64,765 |
| 4 | 9,017 | 10,470 | 45,066 | 1,174 | 64,553 |
| 5 | 9,595 | 11,379 | 43,623 | 1,130 | 64,597 |
| 6 | 11,176 | 11,204 | 41,567 | 1,780 | 63,947 |

**Supplementary Table 3.** QTL-Seq analysis results for Bulk 3/4, showing QTL peaks passing the 99% confidence threshold and supported by >3 SNPs.

| scaffold | start | end | length | nSNPs | avgSNPs_Mb | peakDeltaSNP | posPeakDeltaSNP | avgDeltaSNP |
|---|---|---|---|---|---|---|---|---|
| JN3_SC03 | 90652 | 91482 | 830 | 19 | 22892 | -0.755581831 | 91482 | -0.755094765 |
| JN3_SC03 | 95429 | 95510 | 81 | 4 | 49383 | -0.762071399 | 95510 | -0.762016219 |
| JN3_SC03 | 117995 | 119481 | 1486 | 4 | 2692 | -0.77721984 | 117995 | -0.775634911 |
| JN3_SC03 | 120262 | 120397 | 135 | 4 | 29630 | -0.772987762 | 120262 | -0.772886021 |
| JN3_SC03 | 120490 | 120903 | 413 | 6 | 14528 | -0.772562128 | 120490 | -0.772196231 |
| JN3_SC03 | 121269 | 121376 | 107 | 4 | 37383 | -0.771107876 | 121269 | -0.771046271 |
| JN3_SC03 | 121587 | 123919 | 2332 | 12 | 5146 | -0.770514228 | 121587 | -0.768922143 |
| JN3_SC03 | 129177 | 129485 | 308 | 4 | 12987 | -0.756345077 | 129177 | -0.75613646 |
| JN3_SC03 | 129797 | 130947 | 1150 | 7 | 6087 | -0.75518765 | 129797 | -0.754668941 |
| JN3_SC03 | 133315 | 133575 | 260 | 5 | 19231 | -0.748620183 | 133315 | -0.748391311 |
| JN3_SC03 | 143575 | 144402 | 827 | 6 | 7255 | -0.695954643 | 143575 | -0.693746284 |
| JN3_SC03 | 214154 | 214937 | 783 | 7 | 8940 | -0.787454851 | 214937 | -0.785880757 |
| JN3_SC03 | 254133 | 254842 | 709 | 4 | 5642 | -0.816899703 | 254133 | -0.816126807 |
| JN3_SC03 | 256785 | 257014 | 229 | 4 | 17467 | -0.811341141 | 256785 | -0.811014167 |
| JN3_SC03 | 257115 | 257429 | 314 | 5 | 15924 | -0.810649465 | 257115 | -0.809955842 |
| JN3_SC03 | 257437 | 257645 | 208 | 7 | 33654 | -0.809711568 | 257437 | -0.8094006 |

| | | | | | | | | |
|---|---|---|---|---|---|---|---|---|
| JN3_SC03 | 257974 | 258671 | 697 | 10 | 14347 | -0.807832258 | 257974 | -0.80643415 |
| JN3_SC03 | 258889 | 259459 | 570 | 12 | 21053 | -0.804630083 | 258889 | -0.803186187 |
| JN3_SC03 | 259477 | 259766 | 289 | 28 | 96886 | -0.802572291 | 259477 | -0.802028596 |
| JN3_SC03 | 259825 | 259974 | 149 | 13 | 87248 | -0.801354414 | 259825 | -0.801126937 |
| JN3_SC03 | 260024 | 260135 | 111 | 6 | 54054 | -0.800657985 | 260024 | -0.800465504 |
| JN3_SC03 | 262397 | 262894 | 497 | 4 | 8048 | -0.792353325 | 262397 | -0.79156503 |
| JN3_SC03 | 263902 | 265397 | 1495 | 8 | 5351 | -0.787086359 | 263902 | -0.783510158 |
| JN3_SC03 | 280475 | 280850 | 375 | 8 | 21333 | -0.729086731 | 280475 | -0.728254252 |
| JN3_SC03 | 286079 | 286206 | 127 | 5 | 39370 | -0.684173246 | 286079 | -0.683664536 |
| JN3_SC03 | 287593 | 288962 | 1369 | 22 | 16070 | -0.670565714 | 287593 | -0.662807198 |
| JN3_SC03 | 289965 | 290480 | 515 | 4 | 7767 | -0.649246648 | 289965 | -0.645813308 |
| JN3_SC03 | 309432 | 309547 | 115 | 10 | 86957 | -0.473925994 | 309432 | -0.47364534 |
| JN3_SC03 | 312121 | 312156 | 35 | 5 | 142857 | -0.449502744 | 312121 | -0.449348339 |
| JN3_SC03 | 340008 | 340303 | 295 | 4 | 13559 | -0.299681799 | 340303 | -0.299465264 |
| JN3_BAC01 | 7103 | 20602 | 13499 | 15 | 1111 | -0.794766171 | 20602 | -0.794744975 |

**Supplementary Table 4.** Merged QTL-Seq analysis results for all bulked progeny. Neighboring QTL within 30kb were merged.

| Bulk | Chromosome | QTL number | Start | End |
|---|---|---|---|---|
| B1/2 | JN3_SC04 | 1 | 1092535 | 1105385 |
| | JN3_SC06 | 1 | 1545172 | 1583893 |
| | JN3_SC06 | 2 | 1738694 | 1756847 |
| | JN3_SC07 | 1 | 261543 | 266501 |
| | JN3_SC09 | 1 | 1053225 | 1061275 |
| | JN3_SC09 | 2 | 1180317 | 1302819 |
| | JN3_SC09 | 3 | 1437410 | 1547856 |
| | JN3_SC09 | 4 | 1606032 | 1611796 |
| | JN3_SC09 | 5 | 1661116 | 1785271 |
| | JN3_SC09 | 6 | 1868892 | 1868892 |
| | JN3_SC11 | 7 | 235663 | 238393 |
| | JN3_SC11 | 8 | 290068 | 412461 |
| | JN3_SC11 | 9 | 1085465 | 1096817 |
| B3/4 | JN3_SC03 | 1 | 62390 | 345050 |
| | JN3_SC13 | 2 | 748253 | 748253 |
| | JN3_BAC01 | 3 | 7103 | 20602 |
| B5/6 | JN3_SC03 | 1 | 62390 | 313512 |
| | JN3_SC05 | 1 | 2032604 | 2052005 |
| | JN3_SC07 | 1 | 2236972 | 2261400 |
| | JN3_SC07 | 2 | 2351112 | 2397750 |
| | JN3_SC09 | 1 | 1452050 | 1523178 |
| | JN3_SC09 | 2 | 1681339 | 1787005 |
| | JN3_SC11 | 1 | 195625 | 428764 |
| | JN3_BAC01 | 1 | 6840 | 20602 |

**Supplementary Table 5.** QTL-Seq analysis results for Bulk 1/2, showing QTL peaks passing the 99% confidence threshold and supported by >3 SNPs.

| scaffold | start | end | length | nSNPs | avgSNPs_Mb | peakDeltaSNP | posPeakDeltaSNP | avgDeltaSNP |
|---|---|---|---|---|---|---|---|---|
| JN3_SC04 | 1092535 | 1098768 | 6233 | 6 | 963 | -0.32821503 | 1097401 | -0.32581162 |
| JN3_SC04 | 1103717 | 1105385 | 1668 | 15 | 8993 | -0.32003312 | 1103717 | -0.31946288 |
| JN3_SC06 | 1545172 | 1583893 | 38721 | 83 | 2144 | 0.38697193 | 1561817 | 0.37369769 |
| JN3_SC07 | 261543 | 266501 | 4958 | 8 | 1614 | -0.29788765 | 266501 | -0.29725419 |
| JN3_SC09 | 1053225 | 1061275 | 8050 | 8 | 994 | 0.31740274 | 1057149 | 0.30664662 |
| JN3_SC09 | 1204228 | 1227170 | 22942 | 26 | 1133 | -0.3804147 | 1204228 | -0.35716364 |
| JN3_SC09 | 1238241 | 1253574 | 15333 | 40 | 2609 | -0.33894706 | 1238241 | -0.3289004 |
| JN3_SC09 | 1283617 | 1302819 | 19202 | 12 | 625 | -0.34945828 | 1288802 | -0.34255259 |
| JN3_SC09 | 1452050 | 1456543 | 4493 | 5 | 1113 | 0.76459198 | 1456543 | 0.73626564 |
| JN3_SC09 | 1465687 | 1481733 | 16046 | 334 | 20815 | 0.78981151 | 1481733 | 0.78031216 |
| JN3_SC09 | 1500112 | 1502106 | 1994 | 8 | 4012 | 0.65488916 | 1500112 | 0.64270998 |
| JN3_SC09 | 1503510 | 1504104 | 594 | 5 | 8418 | 0.61735614 | 1503510 | 0.61480681 |
| JN3_SC09 | 1610946 | 1611796 | 850 | 4 | 4706 | -0.37282962 | 1610946 | -0.36645238 |
| JN3_SC09 | 1663429 | 1674868 | 11439 | 5 | 437 | -0.44883742 | 1674868 | -0.40604233 |
| JN3_SC09 | 1682766 | 1684265 | 1499 | 6 | 4003 | -0.5074763 | 1684265 | -0.5014795 |
| JN3_SC09 | 1692451 | 1697605 | 5154 | 9 | 1746 | -0.50857148 | 1692451 | -0.50572646 |
| JN3_SC09 | 1704207 | 1713025 | 8818 | 16 | 1814 | -0.49699533 | 1704207 | -0.49248251 |
| JN3_SC09 | 1733705 | 1785271 | 51566 | 40 | 776 | -0.48250842 | 1733705 | -0.41666603 |
| JN3_SC11 | 1085465 | 1096817 | 11352 | 4 | 352 | -0.30745558 | 1085971 | -0.30488893 |
| JN3_SC11 | 235663 | 238393 | 2730 | 4 | 1465 | -0.29631596 | 238393 | -0.29603571 |
| JN3_SC11 | 290068 | 344372 | 54304 | 41 | 755 | -0.35576269 | 344372 | -0.32411168 |
| JN3_SC11 | 361855 | 412461 | 50606 | 6 | 119 | -0.37347941 | 361855 | -0.34049102 |

**Supplementary Table 6.** QTL-Seq analysis results for Bulk 5/6, showing QTL peaks passing the 99% confidence threshold and supported by >3 SNPs.

| scaffold | start | end | length | nSNPs | avgSNPs_Mb | peakDeltaSNP | posPeakDeltaSNP | avgDeltaSNP |
|---|---|---|---|---|---|---|---|---|
| JN3_SC03 | 90290 | 91482 | 1192 | 15 | 12584 | -0.872599555 | 91482 | -0.872173821 |
| JN3_SC03 | 95429 | 95510 | 81 | 4 | 49383 | -0.878820316 | 95510 | -0.878767421 |
| JN3_SC03 | 117995 | 119516 | 1521 | 10 | 6575 | -0.902461607 | 117995 | -0.902218411 |
| JN3_SC03 | 120021 | 120397 | 376 | 5 | 13298 | -0.901885062 | 120021 | -0.901817789 |
| JN3_SC03 | 120457 | 120903 | 446 | 7 | 15695 | -0.901760988 | 120457 | -0.901705131 |
| JN3_SC03 | 121269 | 121376 | 107 | 4 | 37383 | -0.901529915 | 121269 | -0.901520524 |
| JN3_SC03 | 121587 | 123919 | 2332 | 10 | 4288 | -0.901439421 | 121587 | -0.901224027 |
| JN3_SC03 | 128171 | 128692 | 521 | 11 | 21113 | -0.899565791 | 128171 | -0.899477677 |
| JN3_SC03 | 129141 | 129485 | 344 | 5 | 14535 | -0.899289755 | 129141 | -0.899256119 |
| JN3_SC03 | 129760 | 130947 | 1187 | 12 | 10110 | -0.899113605 | 129760 | -0.899009356 |
| JN3_SC03 | 133255 | 133729 | 474 | 7 | 14768 | -0.898119022 | 133255 | -0.898062636 |
| JN3_SC03 | 143575 | 144072 | 497 | 5 | 10060 | -0.86921785 | 143575 | -0.868272792 |
| JN3_SC03 | 203146 | 203698 | 552 | 5 | 9058 | -0.749515713 | 203698 | -0.748330269 |
| JN3_SC03 | 214154 | 214937 | 783 | 7 | 8940 | -0.790618594 | 214937 | -0.789149801 |
| JN3_SC03 | 254133 | 254842 | 709 | 4 | 5642 | -0.807634066 | 254133 | -0.8067301 |
| JN3_SC03 | 256785 | 257014 | 229 | 4 | 17467 | -0.801132866 | 256785 | -0.800750443 |
| JN3_SC03 | 257115 | 257645 | 530 | 12 | 22642 | -0.800323894 | 257115 | -0.799207878 |

| | | | | | | | | |
|---|---|---|---|---|---|---|---|---|
| JN3_SC03 | 257974 | 258671 | 697 | 10 | 14347 | -0.79727632 | 257974 | -0.795777553 |
| JN3_SC03 | 258889 | 259459 | 570 | 12 | 21053 | -0.7938436 | 258889 | -0.79229575 |
| JN3_SC03 | 259477 | 259766 | 289 | 28 | 96886 | -0.791637655 | 259477 | -0.791054816 |
| JN3_SC03 | 259825 | 259974 | 149 | 13 | 87248 | -0.790332096 | 259825 | -0.790088242 |
| JN3_SC03 | 260024 | 260135 | 111 | 6 | 54054 | -0.789585526 | 260024 | -0.789379188 |
| JN3_SC03 | 262397 | 262894 | 497 | 4 | 8048 | -0.780682964 | 262397 | -0.779837915 |
| JN3_SC03 | 263496 | 265333 | 1837 | 8 | 4355 | -0.776559948 | 263496 | -0.772094599 |
| JN3_SC03 | 280475 | 280859 | 384 | 9 | 23438 | -0.712861421 | 280475 | -0.711908096 |
| JN3_SC03 | 284121 | 284297 | 176 | 5 | 28409 | -0.684713999 | 284121 | -0.683647816 |
| JN3_SC03 | 286079 | 286381 | 302 | 7 | 23179 | -0.666686476 | 286079 | -0.665592145 |
| JN3_SC03 | 287324 | 288684 | 1360 | 13 | 9559 | -0.655223622 | 287324 | -0.646556189 |
| JN3_SC03 | 288759 | 288812 | 53 | 4 | 75472 | -0.642011417 | 288759 | -0.64178124 |
| JN3_SC03 | 288870 | 289198 | 328 | 6 | 18293 | -0.640989428 | 288870 | -0.63981859 |
| JN3_SC03 | 289965 | 290480 | 515 | 4 | 7767 | -0.630907641 | 289965 | -0.627390525 |
| JN3_SC03 | 309432 | 309464 | 32 | 7 | 218750 | -0.459322458 | 309432 | -0.459198624 |
| JN3_SC03 | 312121 | 312315 | 194 | 6 | 30928 | -0.440058592 | 312121 | -0.439725468 |
| JN3_SC05 | 2032604 | 2052005 | 19401 | 28 | 1443 | 0.430753321 | 2039773 | 0.430180259 |
| JN3_SC07 | 2259697 | 2261400 | 1703 | 4 | 2349 | 0.439011021 | 2259697 | 0.437307766 |
| JN3_SC07 | 2362595 | 2366740 | 4145 | 5 | 1206 | -0.471941543 | 2366740 | -0.471341215 |

| | | | | | | | | |
|---|---|---|---|---|---|---|---|---|
| JN3_SC09 | 1452050 | 1456543 | 4493 | 5 | 1113 | 0.829722161 | 1456543 | 0.800513259 |
| JN3_SC09 | 1466929 | 1481733 | 14804 | 291 | 19657 | 0.867284921 | 1481733 | 0.853296369 |
| JN3_SC09 | 1500272 | 1502106 | 1834 | 12 | 6543 | 0.774773349 | 1500272 | 0.766055638 |
| JN3_SC09 | 1502468 | 1504104 | 1636 | 12 | 7335 | 0.757649831 | 1502468 | 0.75051958 |
| JN3_SC09 | 1681339 | 1684265 | 2926 | 16 | 5468 | -0.47546806 | 1684265 | -0.46790509 |
| JN3_SC09 | 1691396 | 1719799 | 28403 | 45 | 1584 | -0.47690688 | 1691396 | -0.468564972 |
| JN3_SC09 | 1733078 | 1787005 | 53927 | 57 | 1057 | -0.514684552 | 1751398 | -0.480106915 |
| JN3_SC11 | 195625 | 202721 | 7096 | 5 | 705 | -0.46833743 | 202721 | -0.464760958 |
| JN3_SC11 | 222987 | 227472 | 4485 | 12 | 2676 | -0.476122405 | 222987 | -0.476028801 |
| JN3_SC11 | 230194 | 230720 | 526 | 4 | 7605 | -0.475785606 | 230194 | -0.475771902 |
| JN3_SC11 | 232822 | 248728 | 15906 | 22 | 1383 | -0.475662794 | 232822 | -0.474713612 |
| JN3_SC11 | 253825 | 317243 | 63418 | 40 | 631 | -0.472159171 | 253825 | -0.452557457 |
| JN3_SC11 | 335201 | 428764 | 93563 | 17 | 182 | -0.489609938 | 397291 | -0.46300473 |
| JN3_BAC01 | 6840 | 20602 | 13762 | 19 | 1381 | -0.890684607 | 6840 | -0.889996515 |

**Supplementary Table 7.** AT-rich (R0) and GC-equilibrated (R1) regions on JN3 scaffold 9, identified by OcculterCut.

| **AT-type** | **Start** | **End** | **ID** |
|---|---|---|---|
| R0 | 1 | 16083 | 0 |
| R1 | 16084 | 30767 | 0 |
| R0 | 30768 | 63011 | 1 |
| R1 | 63012 | 66796 | 1 |
| R0 | 66797 | 69176 | 2 |
| R1 | 69177 | 95332 | 2 |
| R0 | 95333 | 97390 | 3 |
| R1 | 97391 | 135188 | 3 |
| R0 | 135189 | 312804 | 4 |
| R1 | 312805 | 346585 | 4 |
| R0 | 346586 | 349880 | 5 |
| R1 | 349881 | 419334 | 5 |
| R0 | 419335 | 422780 | 6 |
| R1 | 422781 | 585945 | 6 |
| R0 | 585946 | 596083 | 7 |
| R1 | 596084 | 636504 | 7 |
| R0 | 636505 | 656774 | 8 |
| R1 | 656775 | 749330 | 8 |
| R0 | 749331 | 793166 | 9 |
| R1 | 793167 | 805667 | 9 |
| R0 | 805668 | 849060 | 10 |
| R1 | 849061 | 860367 | 10 |
| R0 | 860368 | 875320 | 11 |
| R1 | 875321 | 880003 | 11 |
| R0 | 880004 | 895163 | 12 |
| R1 | 895164 | 898507 | 12 |
| R0 | 898508 | 900486 | 13 |
| R1 | 900487 | 973263 | 13 |
| R0 | 973264 | 986216 | 14 |
| R1 | 986217 | 1094058 | 14 |
| R0 | 1094059 | 1141081 | 15 |
| R1 | 1141082 | 1392212 | 15 |
| R0 | 1392213 | 1415023 | 16 |
| R1 | 1415024 | 1449645 | 16 |
| R0 | 1449646 | 1461842 | 17 |

| | | | |
|---|---|---|---|
| R1 | 1461843 | 1484346 | 17 |
| R0 | 1484347 | 1485396 | 18 |
| R1 | 1485397 | 1486456 | 18 |
| R0 | 1486457 | 1501026 | 19 |
| R1 | 1501027 | 1533064 | 19 |
| R0 | 1533065 | 1818564 | 20 |
| R1 | 1818565 | 1821045 | 20 |
| R0 | 1821046 | 1833561 | 21 |
| R1 | 1833562 | 1857051 | 21 |
| R0 | 1857052 | 1865711 | 22 |
| R1 | 1865712 | 1884012 | 22 |
| R0 | 1884013 | 1889711 | 23 |
| R1 | 1889712 | 2012716 | 23 |
| R0 | 2012717 | 2015980 | 24 |
| R1 | 2015981 | 2025210 | 24 |
| R0 | 2025211 | 2121901 | 25 |

**Supplementary Table 8.** Allelic variation at the *AvrLep2* locus in a collection of 37 isolates virulent or avirulent towards *LepR2* and polymorphic sites in its protein.

| | | | | | Amino acid position | | |
|---|---|---|---|---|---|---|---|
| Isolate | Country[a] | Year[b] | *LepR2*[c] | Nuclotide position[d] | 17 | 93 | 95 |
| JN3 | France | n/a | V | $G^{49}$,$G^{277}$, $G^{278}$, $G^{284}$ | Ala | Gly | Arg |
| 3R11 | Australia | n/a | A | $G^{49}$,$A^{277}$, $A^{278}$, $G^{284}$ | Ala | Asn | Arg |
| 99-56 | Canada, MB | 1999 | A | $G^{49}$,$G^{277}$, $A^{278}$, $A^{284}$ | Ala | Asp | Gln |
| 98-15 | Canada, SK | 1998 | A | $G^{49}$,$G^{277}$, $A^{278}$, $A^{284}$ | Ala | Asp | Gln |
| 290 | France | 1985 | V | $G^{49}$,$G^{277}$, $G^{278}$, $G^{284}$ | Ala | Gly | Arg |
| 2354 | Canada, ON | 1989 | A | $G^{49}$,$A^{277}$, $A^{278}$, $G^{284}$ | Ala | Asn | Arg |
| 99-79 | Canada, SK | 1999 | A | $G^{49}$,$G^{277}$, $A^{278}$, $A^{284}$ | Ala | Asp | Gln |
| 7.1 | Canada, AB | 2005 | V | $G^{49}$,$G^{277}$, $G^{278}$, $G^{284}$ | Ala | Gly | Arg |
| 98-16 | Canada, SK | 1998 | A | $G^{49}$,$G^{277}$, $A^{278}$, $A^{284}$ | Ala | Asp | Gln |
| 00-100 | Canada, MB | 2000 | A | $G^{49}$,$G^{277}$, $A^{278}$, $A^{284}$ | Ala | Asp | Gln |
| 89-13 | Canada | 1989 | A | $A^{49}$,$A^{277}$, $A^{278}$, $G^{284}$ | Thr | Asn | Arg |
| 87-41 | USA | 1987 | A | $G^{49}$,$G^{277}$, $A^{278}$, $A^{284}$ | Ala | Asp | Gln |
| WA51 | Australia | 1989 | A | $G^{49}$,$G^{277}$, $A^{278}$, $A^{284}$ | Ala | Asp | Gln |
| WA30 | Australia | 1989 | A | $G^{49}$,$A^{277}$, $A^{278}$, $G^{284}$ | Ala | Asn | Arg |
| Liffolle5 | France | unknown | A | $G^{49}$,$A^{277}$, $A^{278}$, $G^{284}$ | Ala | Asn | Arg |
| Liffolle6 | France | unknown | A | $G^{49}$,$G^{277}$, $A^{278}$, $A^{284}$ | Ala | Asp | Gln |
| SC07-59 | Canada, SK | 2007 | A | $G^{49}$,$G^{277}$, $A^{278}$, $A^{284}$ | Ala | Asp | Gln |
| IH08-10 | Canada, SK | 2008 | A | $G^{49}$,$G^{277}$, $A^{278}$, $A^{284}$ | Ala | Asp | Gln |
| CB07-37 | Canada, MB | 2007 | A | $G^{49}$,$G^{277}$, $A^{278}$, $A^{284}$ | Ala | Asp | Gln |
| SC07-69 | Canada, SK | 2007 | A | $G^{49}$,$G^{277}$, $A^{278}$, $A^{284}$ | Ala | Asp | Gln |
| VR08-01 | Canada, AB | 2008 | A | $G^{49}$,$G^{277}$, $A^{278}$, $A^{284}$ | Ala | Asp | Gln |
| VR08-29 | Canada, AB | 2008 | A | $G^{49}$,$G^{277}$, $A^{278}$, $A^{284}$ | Ala | Asp | Gln |
| IH08-85 | Canada, SK | 2008 | A | $G^{49}$,$G^{277}$, $A^{278}$, $A^{284}$ | Ala | Asp | Gln |
| PC07-12 | Canada, MB | 2007 | A | $G^{49}$,$G^{277}$, $A^{278}$, $A^{284}$ | Ala | Asp | Gln |
| PC07-45 | Canada, MB | 2007 | A | $G^{49}$,$G^{277}$, $A^{278}$, $A^{284}$ | Ala | Asp | Gln |
| 2367 | Canada, ON | 1989 | A | $G^{49}$,$A^{277}$, $A^{278}$, $G^{284}$ | Ala | Asn | Arg |
| 99-22 | Canada, SK | 1999 | A | $G^{49}$,$G^{277}$, $A^{278}$, $A^{284}$ | Ala | Asp | Gln |
| A94 | Canada | n/a | A | $G^{49}$,$G^{277}$, $A^{278}$, $A^{284}$ | Ala | Asp | Gln |
| 86-12 | Canada, MB | 1986 | A | $G^{49}$,$G^{277}$, $A^{278}$, $A^{284}$ | Ala | Asp | Gln |
| CR07-15 | Canada, AB | 2007 | A | $G^{49}$,$G^{277}$, $A^{278}$, $A^{284}$ | Ala | Asp | Gln |
| 165 | Canada | unknown | A | $G^{49}$,$G^{277}$, $A^{278}$, $A^{284}$ | Ala | Asp | Gln |
| 05-29 | Canada, AB | 2005 | A | $G^{49}$,$G^{277}$, $G^{278}$, $G^{284}$ | Ala | Gly | Arg |
| 04-49 | Canada, MB | 2004 | A | $G^{49}$,$G^{277}$, $A^{278}$, $A^{284}$ | Ala | Asp | Gln |
| 05-08 | Canada, AB | 2005 | A | $G^{49}$,$G^{277}$, $G^{278}$, $G^{284}$ | Ala | Gly | Arg |
| 89-21 | Australia | 1989 | A | $G^{49}$,$A^{277}$, $A^{278}$, $G^{284}$ | Ala | Asn | Arg |
| 166 | Canada | unknown | A | $G^{49}$,$G^{277}$, $A^{278}$, $A^{284}$ | Ala | Asp | Gln |
| 03-02 | Canada, MB | 2003 | A | $G^{49}$,$G^{277}$, $A^{278}$, $A^{284}$ | Ala | Asp | Gln |

[a] Country/province where the isolate was collected ; AB – Alberta, MB – Manitoba, ON – Ontario, SK – Saskatchewan

[b] Year isolate was collected

[c] Interaction phenotype determined following cotyledon-inoculation tests on a differential line harboring Topas-*LepR2* (V, virulent isolate; A, avirulent isolate; I, intermeditae)

[d]Ala: Alanine, Gly: Glycine, Arg: Arginine, Asp: Aspartate, GLn: Glutamine, Asn: Asparagine and Thr: Threonine

**Supplementary Table 9**. Cotyledon and adult plant inoculation tests with virulent and avirulent isolates toward *LepR2*.

| Isolate[a] | | | Cotyledon inoculation test | | | | Adult Plant inoculation test | | | | | | | | | | | | |
|---|---|---|---|---|---|---|---|---|---|---|---|---|---|---|---|---|---|---|---|
| | Location | Year | Topas DH16516 | Topas-*LepR2* | 72-1 (*RlmS*) | $N^{278}$ | Line | Plant 1 | Plant 2 | Plant 3 | Plant 4 | Plant 5 | Plant 6 | Plant 7 | Plant 8 | Plant 9 | Plant 10 | Median | P[d] |
| v23.1.3 | France | n/a | avr (9) | avr (9) | avr (7) | G | DHT | 10 | 10 | 10 | 10 | 10 | 10 | 10 | 10 | 10 | 10 | 10 | <0.0001 |
| | | | | | | | T-*LepR2* | 4 | 3 | 5 | 2 | 7 | 9 | 4 | 3 | 2 | 2 | 4 | *** |
| 00-100 | Canada (MB) | 2000 | avr (9) | Avr (3) | Avr (3) | A | DHT | 9 | 2 | 7 | 7 | 8 | 10 | 9 | 7 | 9 | 4 | 7.5 | <0.0001 |
| | | | | | | | T-*LepR2* | 0 | 0 | 1 | 1 | 0 | 1 | 0 | 0 | 0 | 1 | 0 | *** |
| B15-10 | Canada (SK) | 2015 | avr (9) | avr (7) | avr (8) | A | DHT | 10 | 10 | 9 | 9 | 9 | 5 | 9 | 8 | 10 | 9 | 9 | <0.0001 |
| | | | | | | | T-*LepR2* | 0 | 2 | 0 | 0 | 0 | 1 | 0 | 2 | 0 | 1 | 0 | *** |
| B16-14 | Canada (SK) | 2016 | avr (9) | avr (7) | avr (9) | G | DHT | 8 | 9 | 10 | 10 | 10 | 10 | 10 | 10 | 10 | 10 | 10 | <0.0001 |
| | | | | | | | T-*LepR2* | 1 | 0 | 0 | 1 | 3 | 0 | 3 | 2 | 3 | 1 | 1 | *** |
| B17-12 | Canada (SK) | 2017 | avr (9) | avr (9) | avr (8) | G | DHT | 9 | 10 | 10 | 10 | 10 | 10 | 10 | 10 | 10 | 10 | 10 | 0.0002 |
| | | | | | | | T-*LepR2* | 4 | 0 | 3 | 8 | 5 | 3 | 1 | 3 | 2 | 1 | 3 | *** |
| B17-17 | Canada (SK) | 2017 | avr (9) | avr (9) | avr (7) | G | DHT | 10 | 9 | 8 | 6 | 10 | 10 | 10 | 10 | 10 | 10 | 10 | 0.0004 |
| | | | | | | | T-*LepR2* | 10 | 3 | 5 | 2 | 2 | 1 | 3 | 0 | 2 | 1 | 2 | *** |
| B16-13 | Canada (SK) | 2016 | avr (9) | avr (7) | avr (9) | x | DHT | 1 | 9 | 9 | 10 | 9 | 7 | 10 | 8 | 7 | 4 | 8.5 | 0.603 |
| | | | | | | | T-*LepR2* | 4 | 6 | 10 | NA | 6 | 8 | 7 | 10 | 9 | 3 | 7.0 | NS |
| B18-10 | Canada (SK) | 2018 | avr (9) | avr (9) | avr (9) | x | DHT | 10 | 10 | 10 | 10 | 10 | 10 | 10 | 10 | 10 | 10 | 10 | 0.215 |
| | | | | | | | T-*LepR2* | 10 | 10 | 10 | 9 | 10 | 9 | 10 | 10 | 10 | 9 | 10 | NS |
| B18-11 | Canada (SK) | 2018 | avr (9) | avr (9) | avr (9) | x | DHT | 10 | 9 | 10 | 10 | 8 | 10 | 10 | 10 | 8 | 2 | 10 | 0.087 |
| | | | | | | | T-*LepR2* | 10 | 10 | 10 | 10 | 10 | 10 | 10 | 10 | 10 | 10 | 10 | NS |
| AI397 | Chile | 2005 | avr (9) | avr (7) | avr (7) | A | DHT | 10 | 10 | 10 | 10 | 10 | 10 | 10 | 10 | 10 | 10 | 10 | 0.0.28 |
| | | | | | | | T-*LepR2* | 9 | 10 | 10 | 10 | 10 | 10 | 9 | 3 | 9 | 6 | 9.5 | * |

[a] *L. maculans* isolates used in the study, the location and year in which they were collected MB = Manitoba, SK = Saskatchewan. v23.1.3 is the progeny of an *in vitro* cross (Rouxel et al., 2011).

[b] Classification of isolates in their interaction with *B. napus* lines carrying no *R* genes (Topas DH16516), *LepR2* (Topas-*LepR2*) or *RlmS* (72-1). Interactions classified as avirulent (Avr, black) or virulent (avr, red). Number in brackets indicates median rating score based on 0-9 scale (Larkan *et al.*, 2013). N278 indicates nucleotide at position 278 of the *AvrLep2* coding region, x = deletion of *AvrLep2*.

[c] Results of adult plant test for each isolate infecting Topas DH16516 (DHT) or Topas-*LepR2* (T-LepR2) (ten plants per line). Numbers indicate internal infection score (0-10) for each individual plant, scored 8-10 weeks post-inoculation. Median rating score of each test is given in final column.

[d] Probability and significance of the Mann-Whitney bilateral test for difference between internal infection on Topas DH16516 and Topas-*LepR2. NS,* non-significant; *, significant differences at $p < 0.05$; ***, $p < 0.001$)

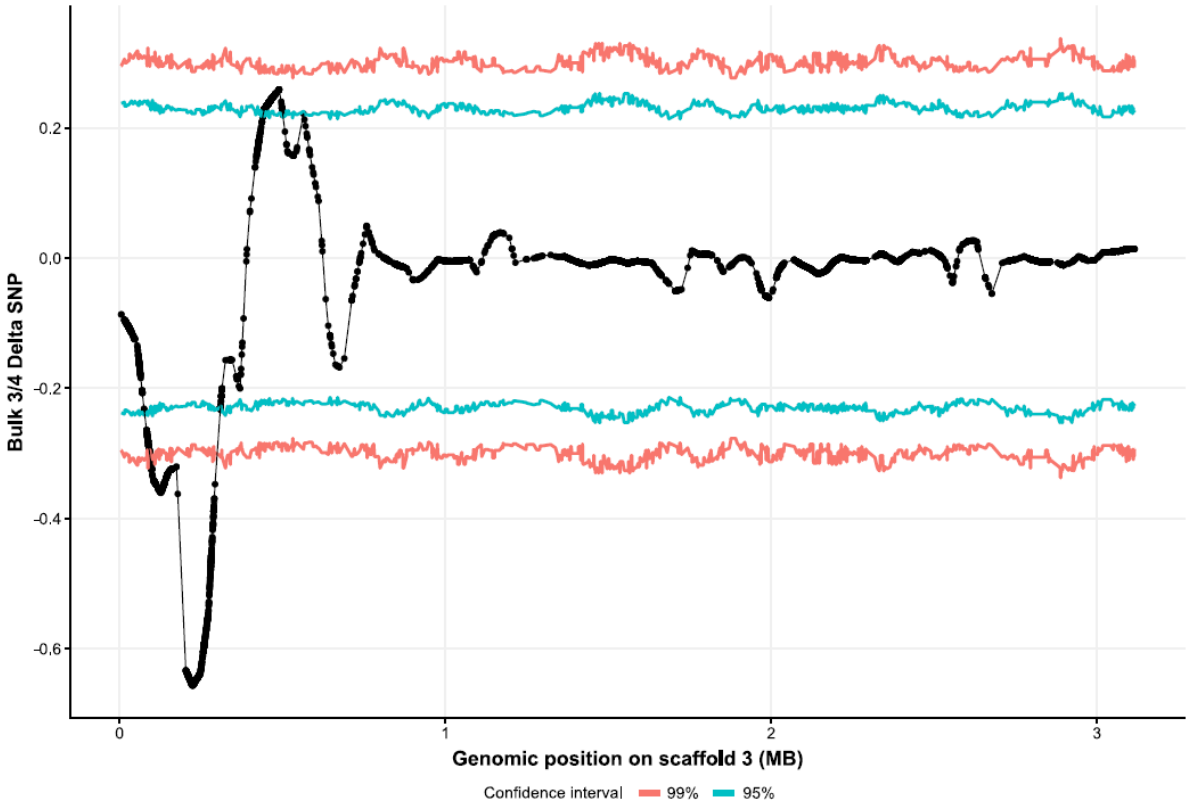


**Supplementary Figure 1.** Plot of Δ(SNP-index) between Bulk 3 and Bulk 4 (*AvrLm7* and *avrLm7*) across scaffold JN3_SC03. Confidence intervals of 95% (blue) and 99% (red) are shown. QTL coordinates are provided in Supplementary Table 3.

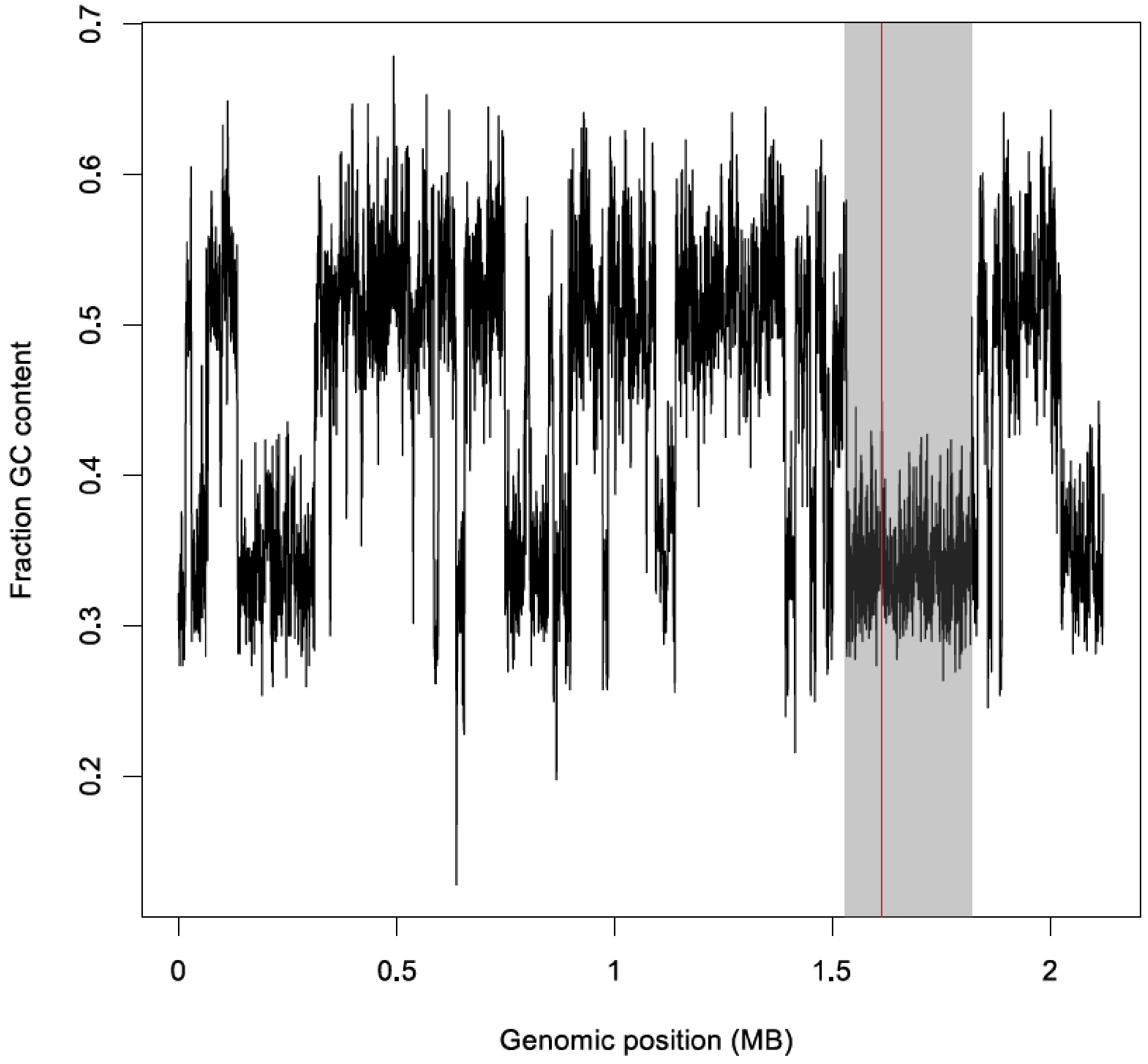


**Supplementary Figure 2.** Plot of GC content calculated in 500 bp windows across scaffold 9. The grey region demarcate the candidate regions identified using QTL-Seq for Bulk 1/2 and Bulk 5/6. The red line shows the CDS of the candidate gene Lmb_jn3_08343.

A

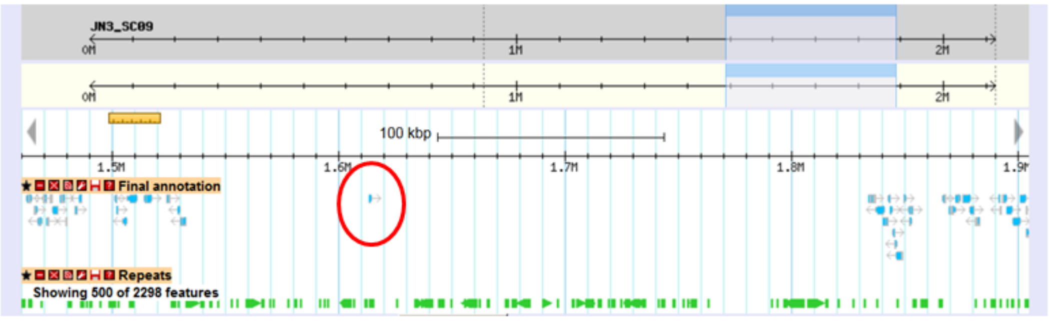


B

>Lmb_jn3_08343 assembled CDS
**MLYLSFYLSLIFFITSATA**TPLFQKKNCHRVVITHADYRSRLTGDSDCSITCQDFLTKDLANFKNRMTQ
IEDNLGGCIRDGSSGVVRETSFIGGRCLCTLKDTLWRMHKACTFEMIMDQYFSLGSDGFGGYRRS
GPISCHA*

**Supplementary Figure 3.** Genomic location and characteristics of the *AvrLmS* (Lmb_jn3_08343) gene in *Leptosphaeria maculans.* **A**, a view of the genomic region hosting Lmb_jn3_08343 (red circle) on the Genoscope genome Browser (http://www.genoscope.cns.fr/leptolifegb/cgi-bin/gbrowse/JN3/); Blue arrows, predicted genes; green arrows, repeated elements. **B**, protein sequence of the gene; the signal peptide is in bold and the cysteine residues in red.

```
  1 ATGCTCTACCTATCTTTCTATTTGTCCTTGATATTCTTTATCACCTCTGCGACTGCAACACCTCTTTTCCAAAAA
  1  M  L  Y  L  S  F  Y  L  S  L  I  F  F  I  T  S  A  T  A  T  P  L  F  Q  K
 76 AAAAACTGTCACAGAGTCGTAATAACGCACGCGGATTACCGATCTCGATTGACTGGCGATTCTGACTGCTCAATA
 26  K  N  C  H  R  V  V  I  T  H  A  D  Y  R  S  R  L  T  G  D  S  D  C  S  I
151 ACATGCCAAGATTTTTAACAAAAGACCTTGCCAATTTTAAGAATCGGATGACACAAATCGAGGATAACCTGGGA
 51  T  C  Q  D  F  L  T  K  D  L  A  N  F  K  N  R  M  T  Q  I  E  D  N  L  G
226 GGCTGTATAAGGGACGGATCGAGCGGTGTCGTCCGAGAGACGTCGTTCATTGATGGACAATGCCTTTGCACCTTA
 76  G  C  I  R  D  G  S  S  G  V  V  R  E  T  S  F  I  D  G  Q  C  L  C  T  L
301 AAAGATACTCTATGGCGAATGCATAAAGCCTGCACTTTTGAAATGATCATGGATCAATATTTCAGCCTTGGATCT
101  K  D  T  L  W  R  M  H  K  A  C  T  F  E  M  I  M  D  Q  Y  F  S  L  G  S
376 GATGGCTTTGGCGGTTATAGGCGGAGCGGTCCCATCAGTTGCCACGCCTAG
126  D  G  F  G  G  Y  R  R  S  G  P  I  S  C  H  A  *
```

A)

```
00-100   AGGGACGGATCGAGCGGTGTCGTCCGAGAGACGTCGTTCATTGATGGACAATGCCTTTGCACCTTA
          R  D  G  S  S  G  V  V  R  E  T  S  F  I  D  G  Q  C  L  C  T  L
v23.1.3  AGGGACGGATCGAGCGGTGTCGTCCGAGAGACGTCGTTCATTGGTGGACGATGCCTTTGCACCTTA
          R  D  G  S  S  G  V  V  R  E  T  S  F  I  Q  G  R  C  L  C  T  L
3R11     AGGGACGGATCGAGCGGTGTCGTCCGAGAGACGTCGTTCATTAATGGACGATGCCTTTGCACCTTA
          R  D  G  S  S  G  V  V  R  E  T  S  F  I  N  G  R  C  L  C  T  L
```

B)

**Supplementary Figure 4.** Nucleotide sequence of the 486 nucleotide region encoding *AvrLep2* from *L.* maculans isolate 00-100 and its predicted amino acid sequence. A) The predicted signal peptide (19 amino acids) and eight cysteine residues are illustrated by gray and green shading respectively. The mutation of codons A278 ->G278 which resulting in a Gly93-> Asp$^{93}$ change in amino acid in the protein leads to the loss of *LepR2*-mediated recognition specificity is indicated in a box. B) Comparison of different haplotypes at *AvrLep2* locus. Non- synonymous base substitutions and their corresponding amino acid changes are indicated by grey shading.

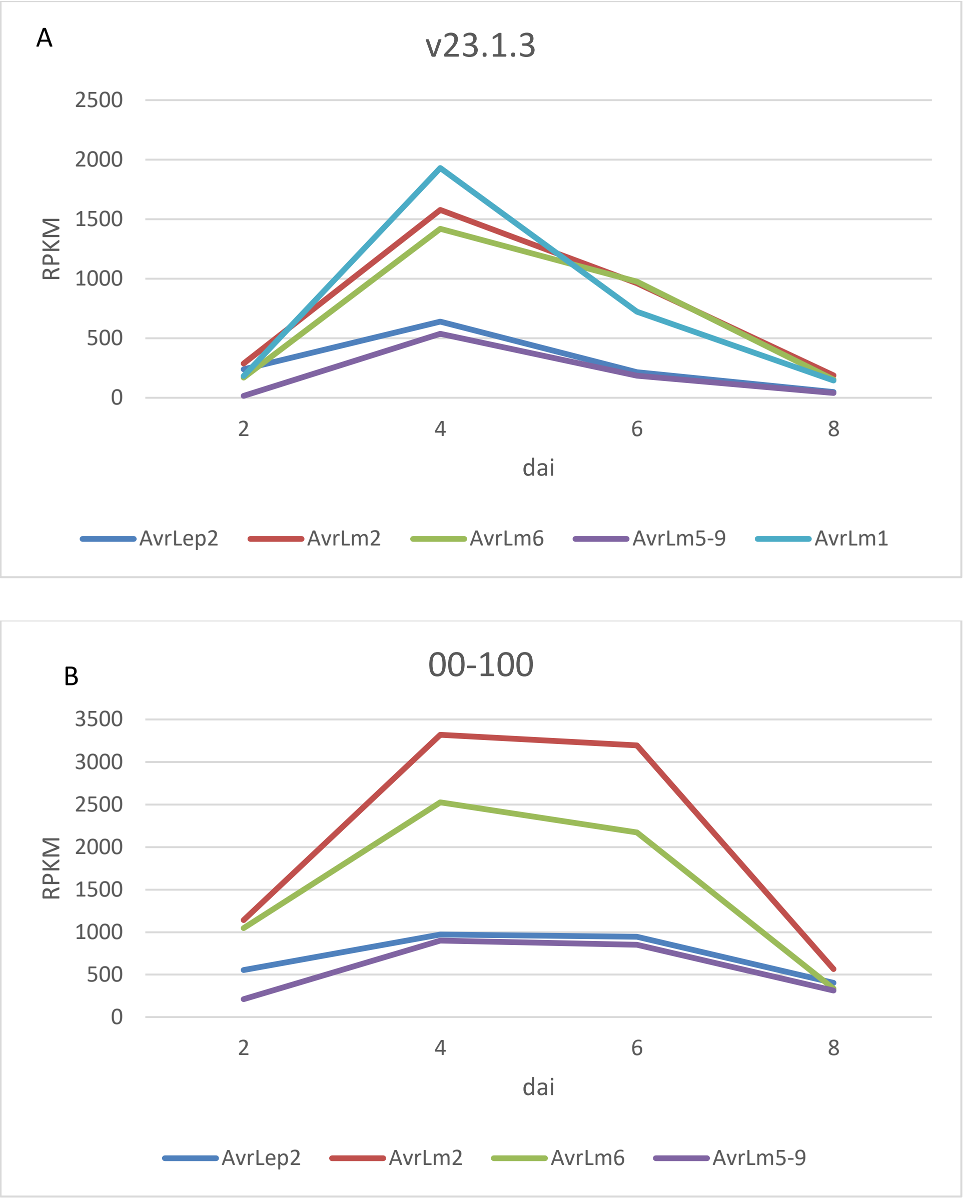


**Supplementary Figure 5**. Comparison of *Avr* gene expression in *L. maculans* isolates v23.1.3 (A) and 00-100 (B) during cotyledon infection of *B. napus* line Topas DH16156. Expression measured as reads per kilobase of transcript per million mapped reads (RPKM) at 2, 4, 6 and 8 days after inoculation (dai).

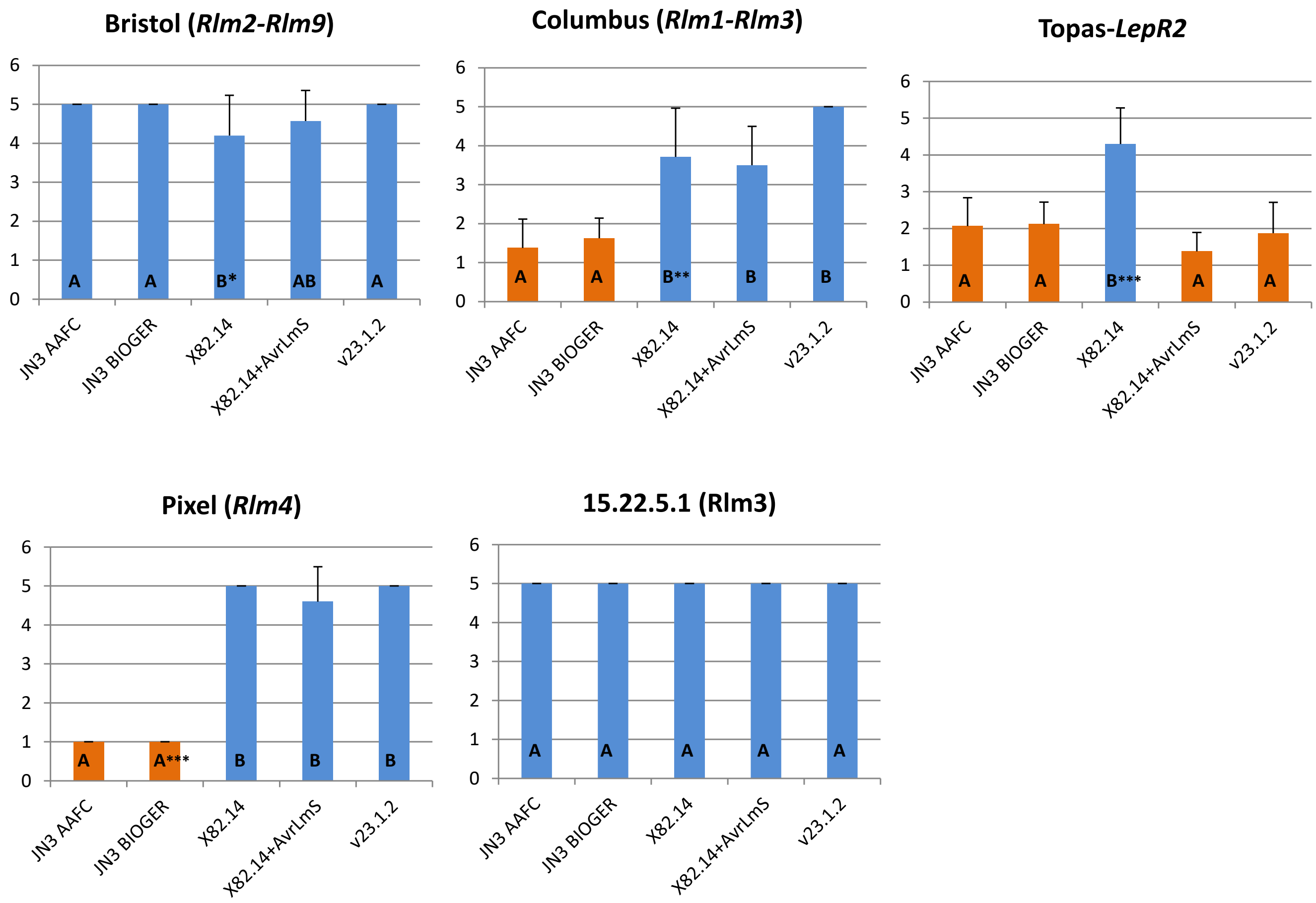


**Supplementary Figure 6.** Comparison of pathogenicity of the two batches of v23.1.3 (JN3) from AAFC and from BIOGER. The two clonal isolates JN3 AAFC and JN3 BIOGER, along with reference isolates used in this study, were inoculated under the protocols and controlled conditions used at BIOGER on the differential plant genotypes Bristol, Columbus, Pixel, 15.22.5.1 and Topas-*LepR2.* The symptoms were rated 16 days post inoculation on a 1 (strong resistance phenotype) to 6 (susceptibility) scale. Values are means and standard deviations of 5-12 inoculation points. Orange bars correspond to avirulent phenotypes (incompatible interactions, mean disease rating <2.5) and blue bars correspond to virulent phenotypes (compatible interactions, mean rating >3.5). A non-parametric Kruskal-Wallis test was performed to detect statistical differences between isolates, which are indicated by different letters on the graphs. *, significant differences at p <0.05; **, p<0.01; ***, p<0.001).
